# Multiple origins of a sex ratio supergene in *Formica ants*

**DOI:** 10.1101/2025.08.27.672632

**Authors:** German Lagunas-Robles, Jessica Purcell, Evan Shimota, Alan Brelsford

## Abstract

Many known supergenes, regions of the genome containing two or more functional mutations with little or no recombination between them, formed in recent evolutionary time to control complex traits. However, when supergenes persist through millions of years of evolution, they may take on additional functions in different lineages. Here, we investigate how an ancient supergene has evolved to gain novel function. In *Formica* ants, most species have an ancient “social” supergene with *M* and *P* haplotypes which determine whether a colony is headed by a single queen or multiple queens, respectively. At least three *Formica* species have an additional haplotype on the same chromosome, termed *M*_*D*_, associated with female-biased offspring sex ratio at the colony level. Is an *M*_*D*_ haplotype present in additional species? Do the *M*_*D*_ haplotypes share a common origin or did they evolve convergently? With whole-genome resequencing, we identify *M*_*D*_ haplotypes in three additional species and use variation across six *Formica* species to examine the evolutionary history of the *M*_*D*_ haplotype. We find that the *M*_*D*_ haplotype did not originate from a rearrangement of the ancestral *M* haplotype. Instead, the *M*_*D*_ haplotype likely originated from recombination between the *M* and *P* haplotypes. Strikingly, we identified two putative origins of the *M*_*D*_ haplotype, one in the ancestor of five Nearctic species, and a second in the Palearctic species *F. cinerea*. The discovery of two *M*_*D*_ haplotypes convergently evolving from distinct recombination events between two ancestral supergene haplotypes illustrates how supergenes can diversify and gain additional phenotypes.

## Introduction

Clusters of linked loci that determine discrete alternative phenotypes are often referred to as “supergenes” (Darlington and Mather 1949, Thompson and Jiggins 2014). Supergenes frequently involve reduced or suppressed recombination, often through structural rearrangements (Schwander et al. 2014). Inversions, where a segment of DNA is reversed in orientation, can preserve associations between alleles at multiple loci on alternative haplotypes. Theoretical work has shown that reduced recombination can maintain beneficial allele combinations (Turner 1967, Jay et al. 2024), but the evolutionary circumstances by which supergenes arise vary from system to system (Schwander et al. 2014, Faria et al. 2019, Gutiérrez-Valencia et al. 2021, Villoutreix et al. 2021, Purcell and Brelsford 2025).

Most described supergenes are found in just one species (e.g., Tuttle et al. 2016, Kim et al. 2017, Funk et al. 2021, Hendrickx et al. 2022, Baird et al. 2023, Jin et al. 2024, Lesturgie et al. 2025). When supergenes are present in multiple congeneric species, they either share a common origin (Brelsford et al. 2020, Yan et al. 2020), or they evolved convergently (Duhamel et al. 2022). Sex chromosomes are a classic supergene example with alternative sexes being determined by alternative haplotypes (Charlesworth 2016). In animal clades with frequent turnover of sex chromosomes, many of the same genes are involved in conserved regulatory networks that result in ovary or testis determination, but the gene that controls which network is switched on and which is switched off can differ between species (Capel 2017). Different genes from within the gonad-determining regulatory networks can take on the role of the switch, but there is a limited pool of such genes. This means that sex chromosomes can evolve independently on homologous genomic regions in multiple species (Jeffries et al. 2018, El Taher et al. 2021). Along the same lines, multiple plant lineages have a supergene that controls flower style length. Emerging consensus suggests that this “S locus” evolved convergently in multiple lineages through distinct gene duplication events (Shang et al. 2025).

While similar genes can lead to the evolution of the same phenotype, convergent supergene phenotypes do not necessarily require the same genes to be present. In some ants, queen number (Purcell et al. 2014, Sigeman et al. 2025) and queen size (Sigeman et al. 2025) are determined by convergent supergenes that do not share gene content. In contrast, rearrangements of similar genomic content led to convergent supergenes in *Microbotryum* mating-type loci (Branco et al. 2018, Duhamel et al. 2022). Remarkably, independent rearrangements occurred at least nine times in this genus (Duhamel et al. 2022), illustrating that the evolution of suppressed recombination can lead to phenotypic convergence. But how do supergenes diversify when they are already established? Does a three-haplotype supergene polymorphism originate through a novel inversion on an established haplotype, or through a recombination event between two divergent haplotypes?

In the ant genus *Formica*, most species have a supergene on chromosome 3 that is associated with colony queen number (Purcell et al. 2014, Brelsford et al. 2020, Lagunas-Robles et al. 2021, McGuire et al. 2022, Pierce et al. 2022, Scarparo et al. 2023, Purcell et al. 2025; but see Sigeman et al. 2024, Lagunas-Robles et al. 2025). Alternative haplotypes determine whether a colony is headed by a single queen (= monogyne) or multiple queens (= polygyne). Monogyne colonies typically have exclusively *M/M* homozygous workers while the *P* haplotype is only found in polygyne colonies. A phylogenetic study of over 90 *Formica* species showed SNPs associated with variation in colony queen number likely predate the ancestor of all extant *Formica* (Purcell et al. 2021), suggesting that the *P* haplotype is an ancient haplotype that arose once.

While the social supergene in *Formica* appears to be ancient, we have more recently discovered additional trait variation that maps to chromosome three in a subset of species. Among the species-specific variation is a haplotype associated with colony sex ratio in three *Formica* species, *F. glacialis, F. podzolica*, and *F. cinerea* (Lagunas-Robles et al. 2021, Scarparo et al. 2023). In *F. glacialis* and *F. podzolica*, monogyne colonies inferred to be headed by a single heterozygous queen (*M*_*D*_/*M*_*A*_) typically produced gynes (= future queens), while monogyne colonies inferred to be headed by a single homozygous queen (*M*_*A*_/*M*_*A*_) produced males (Lagunas-Robles et al. 2021). These two monogyny-associated haplotypes were named after the mythological twins Danaus (*M*_*D*_) and Aegyptus (*M*_*A*_), who respectively had 50 daughters and 50 sons. In *F. cinerea*, monogyne colonies with the *M*_*D*_ haplotype produced gynes more often than monogyne colonies lacking the *M*_*D*_ haplotype (Scarparo et al. 2023). The *M*_*D*_ haplotypes are on chromosome three (Lagunas-Robles et al. 2021, Scarparo et al. 2023), but we do not yet know whether the *M*_*D*_ originated once in a common ancestor of these three species, or convergently in different lineages.

A potential source of genetic variation that may have resulted in the *M*_*D*_ haplotype forming could be the *P* haplotype. The supergene region on chromosome three appears to be dynamic and harbors more than two alternative haplotypes in some species. A “short *P* haplotype” (*P*_*S*_), which spans about half of the extent of the putative ancestral *P* haplotype, was found in *F. glacialis* (Lagunas-Robles et al. 2021) and *F. podzolica* (Purcell et al. 2025). Though it is reduced in size compared to the described *P* haplotypes in other *Formica* species, the *P*_*S*_ haplotype is still associated with polygyny. *Formica podzolica* also has a “long *P* haplotype” (*P*_*L*_; Purcell et al. 2025), which spans the same region of chromosome 3 as the *P* haplotype described in other *Formica* species. In *F. cinerea*, there are two divergent *P* haplotypes, the *P*_1_ and the *P*_2_, both of which span the full supergene region (Scarparo et al. 2023). The recent discovery of a *M* haplotype that contains two *P*-derived genes and is associated with supercoloniality in *Formica paralugubris* and *Formica aquilonia* suggests that novel combinations of two ancient supergene haplotypes can result in the evolution of new phenotypes (Sigeman et al. 2024).

To establish when and how the *M*_*D*_ haplotype evolved, we compared the *M*_*D*_ haplotype in five Nearctic species and one Palearctic species. First, we ask whether there is a single origin or multiple origins of the *M*_*D*_ haplotype. We examined the supergene boundaries of the *M*_*D*_ haplotypes in each species and compared putative haplotype breakpoints. If there was one origin of the *M*_*D*_ haplotype (Figure 1A, hypothesis 1), then all the *M*_*D*_ haplotypes would share haplotype boundaries. If there were multiple origins of the *M*_*D*_ haplotype (Figure 1A, hypothesis 2), then the species with homologous *M*_*D*_ haplotypes could have distinct haplotype boundaries from their non-homologous counterparts. Second, we ask how the *M*_*D*_ haplotype originated – did the *M*_*D*_ haplotype originate from a rearrangement on an ancestral *M* haplotype or did it form through recombination between the *M* and *P* haplotypes? To examine the evolutionary history of the *M*_*D*_ haplotype, we used a windowed phylogenetic approach to scan the chromosome harboring the supergene for regions with a shared history between *M*_*D*_ haplotypes. If the *M*_*D*_ haplotype arose from a rearrangement on the *M* haplotype (Figure 1B, hypothesis 1), then we would expect the *M*_*D*_ haplotype to have more regions with a shared phylogenetic history with the *M*_*A*_ haplotype than the *P* haplotype. If the *M*_*D*_ haplotype arose from recombination between the *M* haplotype and the *P* haplotype (Figure 1B, hypothesis 2), then we would expect *M*_*D*_ haplotypes to have more regions with a shared phylogenetic history with the *P* haplotype. Regions that freely recombine within each species are expected to have topologies matching the species tree and to be concentrated outside the supergene region, a previously documented pattern in species with alternative social supergene haplotypes (Brelsford et al. 2020).

**Figure 1.**
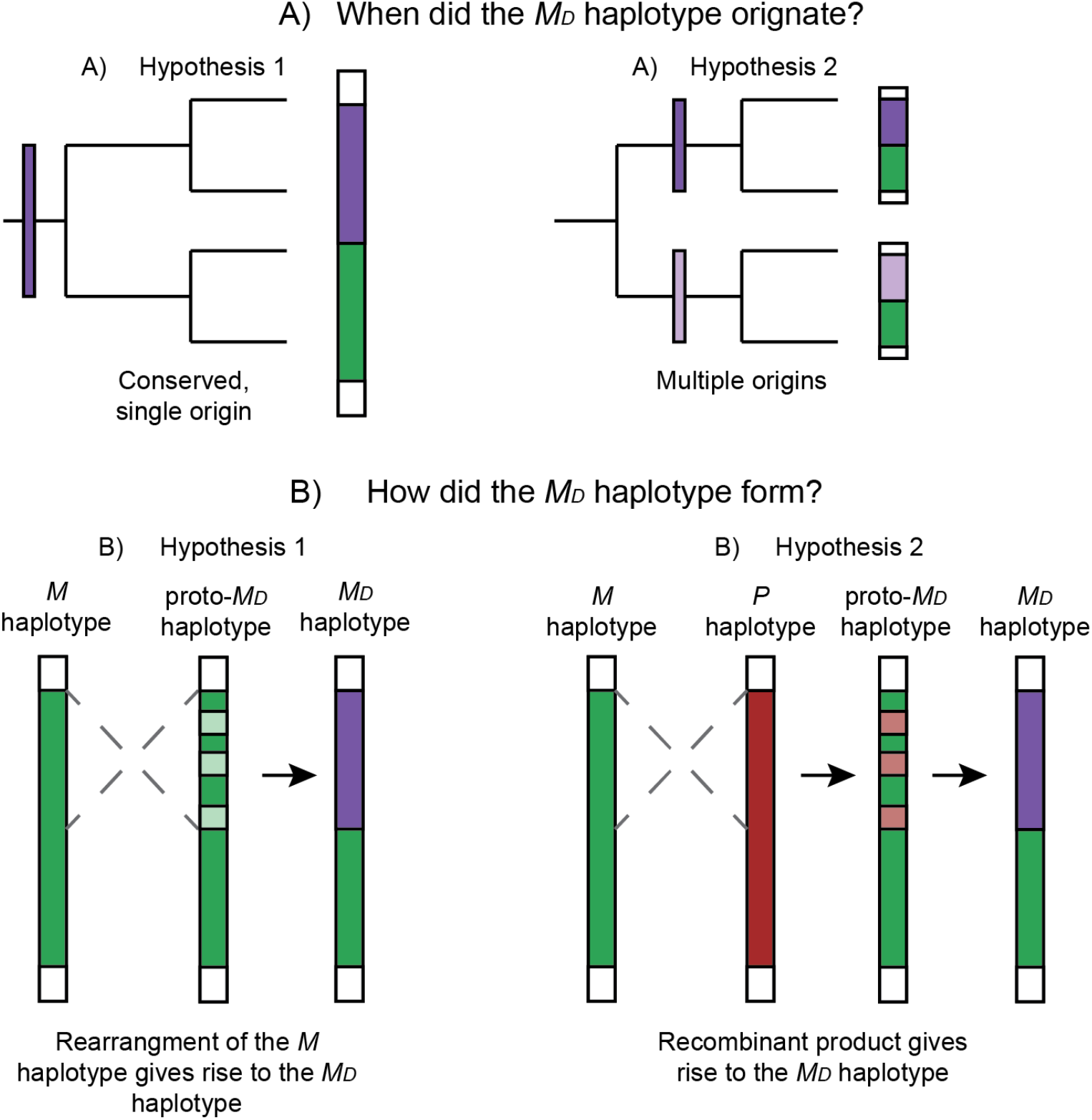
An *M*_*D*_ haplotype is associated with gyne-producing colonies in *F. glacialis, F. podzolica*, and *F. cinerea*. This diagram illustrates the main questions this paper aims to address. **(A)** How many times did the *M*_*D*_ haplotype originate? H1: If the *M*_*D*_ haplotype is conserved and has a single origin, then it would be present in all clades. H2: If the *M*_*D*_ haplotype originated multiple times, then each *M*_*D*_ haplotype variant would be restricted to its respective clade. **(B)** How did the *M*_*D*_ haplotype form? H1: If the *M*_*D*_ haplotype was a rearrangement of the ancestral *M* haplotype, then the *M*_*D*_ haplotype would be more similar to the *M* haplotype than the *P* haplotype across the *M*_*D*_ haplotype region. H2: If the *M*_*D*_ haplotype was a product of a recombination event that placed parts of the ancestral *P* haplotype onto the *M* haplotype background, then the *M*_*D*_ haplotype would be more similar to the *P* haplotype than the *M* haplotype across the recombined region.

## Methods

### Sample collections

In the present study, we included previously sampled species with known *M*_*D*_ haplotypes, two Nearctic species, *F. glacialis* and *F. podzolica* (Lagunas-Robles et al. 2021; Purcell et al. 2025), and a Palearctic species, *F. cinerea* (Scarparo et al. 2023). We sampled three additional Nearctic species, *F. pacifica, F. transmontanis*, and *F*. cf. *transmontanis* (see Table S1), that are closely related to *F. glacialis* and *F. podzolica* (Borowiec et al. 2021). We collected alates (reproductive queens and males) from *F. pacifica* and *F. transmontanis* colonies in August 2022. To determine the association between colony sex ratio and supergene haplotype, we collected at least 5 workers per colony and as many alates as possible. We assessed colony-level sex ratio for colonies with ≥5 alates collected and classified them as male-producing (≥80% males), gyne-producing (≥80% gynes), or mixed.

### DNA extraction and resequencing preparation

To extract genomic DNA, we first ground the ant tissue with a sterilized plastic pestle after immersing the tube in liquid nitrogen and digested the tissue overnight at 56°C in 180µL ATL buffer and 20µL proteinase K. We performed the DNA purifying steps using the QIAamp 96 DNA kit for the QIAcube HT robot following the manufacturer’s protocol. We eluted the samples in 100µL of 10 µM Tris-HCl (pH 8.0). For *F. cinerea*, we used DNA extractions from samples published in Scarparo et al. 2023. We prepared the samples for whole-genome resequencing using the Seqwell plexWell LP-384 library kit following manufacturer’s instructions. The libraries were sequenced on three separate NovaSeq 6000 lanes by Novogene, Inc. using 150 base-pair (bp) paired-end reads. We produced data for 41 *F. glacialis*, 42 *F. pacifica*, 76 *F. transmontanis*, 24 *F*. cf. *transmontanis*, and 43 *F. cinerea*.

### Publicly available data

We used a subset of *M*_*D*_/*M*_*A*_, *M*_*A*_/*M*_*A*_, and *P*_*S*_/*M*_*A*_ *F. glacialis* whole-genome data from Lagunas-Robles et al. 2021 (*n* = 8) and Purcell et al. 2021 (*n* = 2), as well as a subset of *M*_*A*_/*M*_*A*_ and *M*_*D*_/*M*_*A*_ samples for *F. podzolica* from Lagunas-Robles et al. 2021 (*n* = 10) and *M*_*A*_/*M*_*A*_, *M*_*D*_/*M*_*A*_, and *P*_*L*_/*M*_*A*_ samples from Purcell et al. 2025 (*n* = 123). We also used two *F. cinerea* whole-genome resequencing samples from Brelsford et al. 2020 and a single *F*. cf. *transmontanis* from Purcell et al. 2021 (see Table S1 for project accession numbers associated with these samples).

### Data processing

We demultiplexed the raw reads in Stacks version 2.60 using “process_shortreads” (Catchen et al. 2013). We removed adapters and merged overlapping paired-end reads using PEAR version 0.9.11 (Zhang et al. 2014). We then aligned the single and paired reads to the chromosome-level *Formica selysi* genome assembly (Brelsford et al. 2020; NCBI accession GCA_009859135.1) using the with the program bwa-mem2 version 2.2.1 (Vasimuddin et al. 2019). We used “fixmate” and “markdup” to mark and remove PCR duplicates in SAMtools version 1.18 (Li et al. 2009). We then merged and sorted the aligned single and paired read bam files.

### Phylogenetic relationships among the species

In order to establish the species tree topology, we generated a maximum-likelihood tree using whole-genome (WG)-resequencing data. We called variants using bcftools version 1.16 (Li 2011) with “mpileup” and “call” from the BAM files for all species. We filtered the VCF file in VCFtools version 0.1.17 (Danecek et al. 2011) by retaining variants with a minimum mapping quality of 20, and filtered the variants using the following parameters: retained bi-allelic sites (--max-alleles 2), minimum depth of 2 (--minDP 2), minor allele frequency of 0.05 (--maf 0.05), allowed up to 15% missing data per SNP (--max-missing 0.85), and removed indels (--remove-indels). We excluded chromosome 3, the chromosome harboring an ancient *Formica* supergene (Brelsford et al. 2020; Purcell et al. 2021), as trans-species variation on chromosome 3 would confound inference of the species tree (Brelsford et al. 2020). We selected a subset of 10 high-coverage (depth between 5.8 and 24.7) individuals for each putative species group to establish phylogenetic relationships. We refiltered the raw data retaining the 60 high-coverage individuals with the same parameters described above resulting in a dataset of 3,206,482 SNPs.

We converted the filtered VCF file containing these 60 individuals to phylip format using vcf2phylip.py (Ortiz 2019). We then produced a maximum-likelihood (ML) tree in IQ-TREE v2.2.2.6 (Minh et al. 2020) enabling ModelFinder Plus (Kalyaanamoorthy et al. 2017), which selected TVM+F+ASC+R8 as the best model. We specified ascertainment corrections (Lewis 2001), which informs ModelFinder Plus that the alignment does not contain invariant sites. Branch support was calculated using 1,000 ultra-fast bootstrap trees with UFBoot2 (Hoang et al. 2018). We visualized the ML tree in FigTree version 1.4.4 (Rambaut 2012).

We identified clades for *F. glacialis, F. podzolica, F. pacifica*, and *F. cinerea. F. transmontanis* has a disjunct geographic distribution (Francoeur 1973), with one population segment found in inland Washington and British Columbia, and another found along the coast of northern California. Our results showed that this species is paraphyletic; the northern population was sister to *F. pacifica*, while the southern population was sister to the ancestor of *F. pacifica* and the northern population, with 100% bootstrap support (Figure S1). We will refer to the northern clade as *F. transmontanis* and the southern clade as *F*. cf. *transmontanis* in this study. We will refer to the two sister species, *F. glacialis* and *F. podzolica* as the “*glacpodz* clade” and to the species trio of *F. pacifica, F. transmontanis*, and *F*. cf. *transmontanis*, as the “*pactrans* clade” (Figure S1).

### Assigning supergene haplotypes

To determine the supergene variation in each species, we examined the known supergene regions of chromosome 3 in *F. glacialis, F. podzolica, F. pacifica, F. transmontanis, F*. cf. *transmontanis*, and *F. cinerea*. We used ANGSD version 0.937 (Korneliussen et al. 2014), which utilizes genotype likelihoods calculated directly from BAM files to account for uncertainty in low-coverage whole-genome data. To generate a principal component analysis (PCA) for the region harboring the *M*_*D*_ haplotype on chromosome 3 in *F. glacialis* and *F. podzolica* (Lagunas-Robles et al. 2021), we first calculated genotype likelihoods for each species separately with the following parameters: “-doMajorMinor 4, -doMaf 1, -skipTriallelic 1, -minMapQ 25, -minQ 25, -r Scaffold03: 2000000-7500000, -remove_bads 1, -GL 1 -doGlf 2, -setMinDepth 1, -SNP_pval 1e-6, -minInd (50% of samples on a per species basis).”

To examine the presence of *P* haplotypes in our samples, we examined the region of 7.5-12.4Mbp on chromosome 3. The different *P* haplotypes *P*_*L*_, *P*_*S*_, *P*_1_, and *P*_2_ are distinct from the *M* haplotypes across this region (Lagunas-Robles et al. 2021, Scarparo et al. 2023, Purcell et al. 2025) and would separate in PC space from the *M* haplotypes. We generated a genotype-likelihood file for “-r Scaffold03: 750000-1240000” for *F. glacialis, F. podzolica*, and *F. cinerea* and tested for the presence of supergene variation in the *P* haplotype region in *F. pacifica, F. transmontanis*, and *F*. cf. *transmontanis*.

We used the genotype likelihood files to calculate species covariance matrices for the first 5 eigenvectors in PCAngsd version 1.10 (Meisner and Albrechtsen 2018). We calculated the eigenvalues with the “eigen” function in R version 4.2.3 (R Core Team 2023). We identified supergene genotype clusters by leveraging individuals with known genotypes from a *F. glacialis* dataset (Lagunas-Robles et al. 2021). We also confirmed the known genotypes for previously analyzed *F. cinerea* (Brelsford et al. 2020, Scarparo et al. 2023), *F*. cf. *transmontanis* (Purcell et al. 2021), and *F. podzolica* samples (Lagunas-Robles et al. 2021, Purcell et al. 2025).

We inferred the supergene genotypes in *F. pacifica, F. transmontanis*, and *F*. cf. *transmontanis* by estimating the inbreeding coefficient (*F*_IS_) in the supergene region of chromosome 3 with variants identified with bcftools mpileup (see above). We filtered variants for each species separately using the following parameters: retained bi-allelic sites (--max-alleles 2), minimum depth of 2 (--minDP 2), minor allele count of 2 (--mac 2), allowed up to 25% missing data per SNP (--max-missing 0.75), and removed indels (--remove-indels) in VCFtools version 0.1.17 (Danecek et al. 2011). We then estimated the inbreeding coefficient (*F*_IS_) for each individual in each filtered species VCF file with the “--het” flag in VCFtools version 0.1.17 (Danecek et al. 2011). Individuals with a homozygous genotype are expected to have higher *F*_IS_ values in the supergene region relative to individuals with a heterozygous genotype (Scarparo et al. 2023). We used a combination of *F*_IS_ and PC1 for each species to determine whether an individual was homozygous or heterozygous for a genotype.

### Estimating the M_D_ haplotype boundaries

In order to estimate the *M*_*D*_ haplotype boundaries, we calculated genetic differentiation (*F*_ST_) between the *M*_*D*_/*M*_*A*_ and *M*_*A*_/*M*_*A*_ individuals for each species. In ANGSD version 0.937 (Korneliussen et al. 2014), we estimated genotype likelihoods for each species supergene genotype group using the SAMtools model “-GL -1” and estimated the site frequency spectrum (SFS) using “-doSaf 1”. We used sites that met a minimum base and mapping quality of 25 (-minQ 25 and -minMapQ 25) and used “remove_bads” set to 1. The SFS was polarized against the *F. selysi* genome assembly (Brelsford et al. 2020; NCBI accession GCA_009859135.1). Then we used species SFSs to generate a two-population frequency spectrum for intraspecific genotypes and estimated *F*_ST_ across non-overlapping 10kb sliding windows (-win 10000 and - step 10000) using “FST stats2” in ANGSD version 0.937 (Korneliussen et al. 2014). We used the qqman package (Turner 2018) to visualize *F*_ST_ across the genome.

### *Evaluating the Presence of a* P *haplotype*

We calculated *F*_ST_ in ANGSD version 0.937 (Korneliussen et al. 2014), using the methods described in “*Estimating the* M_D_ *haplotype boundaries*,” between the genetic clusters showing the highest variance on PC1 in the 7.5-12.4Mbp region. PCAs are expected to capture the genetic variation that distinguishes the non-recombining *M* and *P* haplotypes. After confirming that PC1 consistently separated *M* haplotypes from *P* haplotypes in the three species with known *P* haplotypes, *F. cinerea* (Brelsford et al. 2020, Scarparo et al. 2023), *F. glacialis* (Lagunas-Robles et al. 2021), and *F. podzolica* (Purcell et al. 2025), we searched for a *P* haplotype in the *pactrans* clade by estimating *F*_ST_ between the two genetic clusters showing the highest separation on PC1 in the 7.5-12.4Mbp region for each of the three species.

### *Identifying* M_D_ *haplotypes in the pactrans clade by PCR-RFLP*

In order to design a SNP assay that differentiated the *M*_*D*_ haplotype from the *M*_*A*_ haplotype in the *pactrans* clade, we identified fixed SNPs on the *M*_*D*_ haplotype of all three species. To do so, we filtered the raw VCF file for each species separately using the following parameters: retained biallelic sites (--max-alleles 2), minimum depth of 1 (--minDP 1), minor allele count of 2 (--mac 2), allowed for 25% missing data per SNP (--max-missing 0.75), and removed indels (--removeindels) in VCFtools version 0.1.17 (Danecek et al. 2011). We then calculated allele frequencies for the *M*_*D*_/*M*_*A*_ and *M*_*A*_/*M*_*A*_ individuals in each species separately using “--freq2” in VCFtools version 0.1.17 (Danecek et al. 2011). We identified SNPs on chromosome 3 with a reference allele frequency between 0.4 and 0.6 in *M*_*D*_/*M*_*A*_ individuals and reference or alternate allele frequency > 0.98 in *M*_*A*_/*M*_*A*_ individuals, as would be expected for a fixed difference between the *M*_*D*_ and *M*_*A*_ haplotypes. We verified a SNP at position 2789194 by extracting the genotypes of individuals that had a minimum depth of 6 (--minDP 6) at this position in 012 format using “--012” in VCFtools version 0.1.17 (Danecek et al. 2011). We validated the called genotypes at position 2789194 by comparing the called genotypes to the assigned supergene genotypes on the PCAs (Figure 2A-2C). We found one mismatch out of 64 total individuals. We visually inspected the mismatched individual’s BAM file with SAMtools version 1.18 using “tview” (Li et al. 2009). We found that this mismatch likely results from a genotyping error, with 10 reads for the *M*_*A*_-specific allele, matching the individual’s supergene genotype inferred from the PCA, and one read for the *M*_*D*_-specific allele, which may result from index-hopping or lab contamination (Table S2).

**Figure 2.**
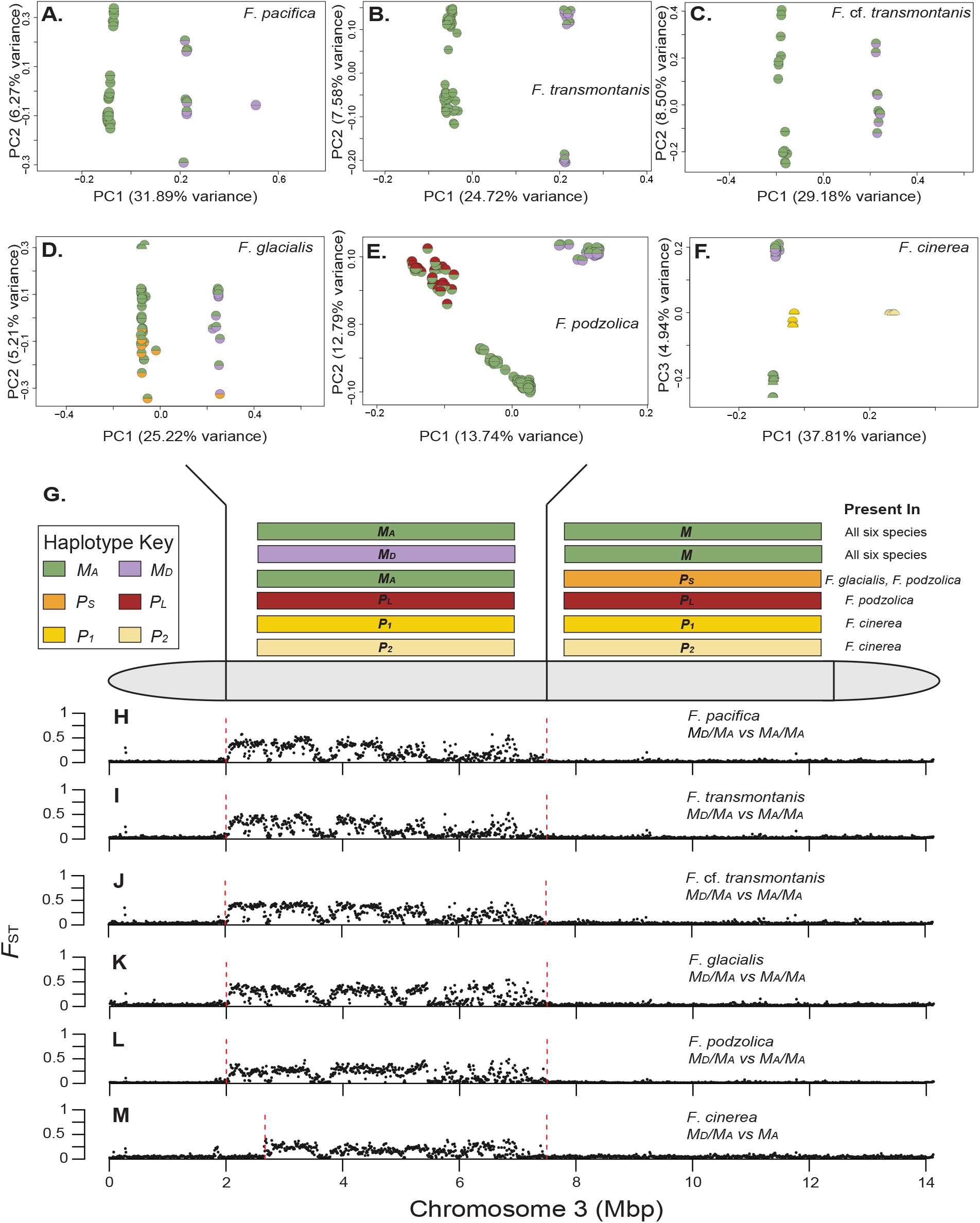
Genotypes were assigned based on principal component analyses by using SNPs on chromosome 3 between 2-7.5Mbp. Each colored half circle represents a haplotype. Since males are haploid, they are represented by half circles. Female worker ants are diploid and are represented by a full circle. The schematic represents where the haplotypes are found on the chromosome. Genetic differentiation (*F*_ST_) is shown between *M*_*D*_/*M*_*A*_ and *M*_*A*_/*M*_*A*_ genotypes. In the 2-7.5Mbp region: **(A)** *F. pacifica* separated into three supergene genotype clusters. PC1 separates individuals into two homozygous clusters on each end (*M*_*A*_/*M*_*A*_ and *M*_*D*_/*M*_*D*_) and the heterozygous cluster (*M*_*D*_/*M*_*A*_) in the middle. **(B and C)** *F. transmontanis* and *F*. cf. *transmontanis* each separated into two supergene genotype clusters along PC1 with geographic population structure shown along PC2. **(D)** *F. glacialis* separated into two supergene genotype clusters along PC1, with *M*_*A*_/*M*_*A*_ and *P*_*S*_/*M*_*A*_ individuals forming one cluster and individuals heterozygous for the *M*_*D*_ haplotype forming the other. **(E)** *F. podzolica* separated into three supergene genotype clusters along PC axes 1 and 2, representing the three genotypes we selected from Purcell et al. 2025 (*M*_*A*_/*M*_*A*_, *M*_*D*_/*M*_*A*_, and *P*_*L*_/*M*_*A*_). **(F)** *F. cinerea* separated into four supergene genotype clusters along PC axes 1 and 2, representing the four genotypes we resequenced from Scarparo et al. 2023 (*M*_*D*_/*M*_*A*_, *M*_*A*,_ *P*_*1*_, and *P*_*2*_). **(G)** A schematic representation of the boundaries of the supergene haplotypes. Genetic differentiation between *M*_*D*_/*M*_*A*_ individuals and *M*_*A*_ homo/hemi-zygotes on chromosome 3 reveals shared *M*_*D*_ haplotype boundaries in the Nearctic species, but a shorter span of the *M*_*D*_ in *F. cinerea*. Each point represents a 10-kbp window. Red lines delimit the estimated boundaries for *M*_*D*_ haplotypes of each species. **(H-J)** *F. pacifica, F. transmontanis*, and *F*. cf. *transmontanis* exhibit elevated *F*_ST_ between 2Mbp and 7.5Mbp. **(K and L)** *F. glacialis* and *F. podzolica* exhibit elevated *F*_ST_ between 2Mbp and 7.5Mbp, a result previously reported in Lagunas-Robles et al. 2021 and Purcell et al. 2025. **(M)** The *M*_*D*_ haplotype in *F. cinerea* has elevated *F*_ST_ between 2.67Mbp and 7.5Mbp.

We designed primers (GGTCCAGGAGATCGCATGT and ATTGACACGACACATGCCG) to amplify a 255 base pair (bp) fragment flanking the haplotype-specific SNP at position 2789194. We performed a PCR at the following temperatures: an initial denaturing step at 94°C for 3 minutes, followed by 35 cycles of denaturing at 94°C for 30 seconds, annealing at 58°C for 30 seconds, and extending at 72°C for 30 seconds, and a final extending step at 72°C for 3 minutes. The PCR amplicon was digested with the restriction enzyme, BspDI, at 37°C for 60 minutes. The enzyme was selected to yield the following restriction fragments: *M*_*A*_/*M*_*A*_ homozygotes result in two fragments at 156 bp and 99 bp, heterozygous *M*_*D*_/*M*_*A*_ individuals result in three fragments at 255 bp, 156 bp, and 99 bp, and *M*_*D*_*/M*_*D*_ homozygotes result in one fragment at 255 bp. We visualized the digested product using 2% agarose gel electrophoresis.

### *The* M_D_ *haplotype function in the pactrans clade*

We examined the genotypes of 145 workers from 28 colonies with sex ratio data. In total, 7 colonies produced gynes, 17 produced males, and 4 produced both gynes and males. For two colonies with known sex ratio, workers (*n* = 5 and *n* = 4, respectively) were included in our WGS dataset. For the remaining 26 colonies, we used the PCR-RFLP assay to assess the presence or absence of the *M*_*D*_ haplotype. Out of the 28 colonies, we collected five alates from two colonies, while the remaining 26 colonies had ≥ 7 alates.

To test for an association between gyne production and the *M*_*D*_ haplotype, we performed a Fisher’s exact test (Fisher 1934) using “fisher.test” in R version 4.2.3 (R Core Team 2023). We generated a contingency table with the following binary colony classifications: For *M*_*D*_ haplotype presence, colonies containing one or more workers with an *M*_*D*_ haplotype were coded as 1, while colonies lacking the *M*_*D*_ haplotype were coded as 0. For presence of gynes, colonies classified as gyne-producing or mixed were coded as 1, while colonies classified as male-producing were coded as 0 (Table S3).

### *Evolutionary history of the* M_D_ *haplotype*

We examined the evolutionary history of supergene haplotypes across species using windowed phylogenies on chromosome 3. Windowed phylogenies allow us to explore the relationships between the different haplotypes across the different parts of the chromosome and identify regions that are conserved between various groups of supergene haplotypes within or across species. Since the *M*_*D*_ haplotype and *P*_*L*_ haplotype are only found in the heterozygous state in diploid females, we leveraged the homozygous *M*_*A*_/*M*_*A*_ with the heterozygous *M*_*D*_/*M*_*A*_ and the heterozygous *P*_*L*_/*M*_*A*_ genotypes to infer haplotype-specific allele frequencies at each SNP on chromosome 3 for the *M*_*D*_ and *P*_*L*_ haplotypes.

We used the “--freq2” flag in VCFtools version 0.1.17 (Danecek et al. 2011) to calculate within-species haplotype-specific allele frequencies for each genotype (*M*_*A*_*/M*_*A*_, *M*_*D*_*/M*_*A*_, and *P*_*L*_/*M*_*A*_) with the filtered VCF file. After retaining only SNPs that were sequenced in at least one individual for every combination of species x genotype, we retained 373,569 SNPs which resulted in 7,472 50-SNP windows for the analysis. To infer the population-level allele frequency of the *M*_*D*_ and *P*_*L*_ haplotypes, we calculated the frequencies of haplotype-specific alleles with the following formulas:

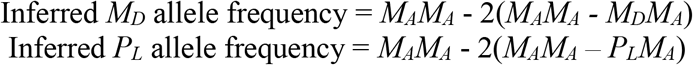

We transformed inferred frequencies that were greater than 1 or negative to 1 or 0, respectively, since these likely represent sampling error. The inferred *M*_*D*_ haplotype allele frequency was calculated for all species in the analysis, while the inferred *P*_*L*_ haplotype allele frequency was calculated for *F. podzolica*. Since we sequenced homozygous *M*_*A*_*/M*_*A*_ workers for all Nearctic species and haploid males for the *F. cinerea M*_*A*_, *P*_*1*_, and *P*_*2*_ haplotypes, we used the reference allele frequencies directly. We generated a table with the allele frequency calculated for each position in the chromosome calculated for each species x haplotype combination in the dataset, some of which were inferred (see above) and some of which were measured directly from haploid males or homozygous females.

We calculated distance matrices using non-overlapping 50-SNP windows with the “dist” function specifying “method = euclidean” in R version 4.2.3 (R Core Team 2023). We used the distance matrices to generate a neighbor-joining tree for each window using “nj” in the R package APE version 5.7.1 (Paradis et al. 2004). We then tested the windowed trees for criteria topologies using “is.monophyletic” in APE. These results were summarized by mapping the SNP windows on chromosome 3 that met each monophyly criterion.

Once we established which 50-SNP windows aligned with each monophyly criterion, we concatenated the windows within each category to produce empirical trees for each of the 11 criterion topologies (Figure S4).

### Read mapping quality on chromosome 3

To examine alignment accuracy near the putative inversion breakpoints for the haplotypes, we used mapping quality, as calculated by bwa-mem2 (Vasimuddin et al. 2019) during read alignment. We selected two individuals for each species with the highest depth for the *M*_*A*_*/M*_*A*_ and *M*_*D*_/*M*_*A*_ genotypes. We selected the highest depth *M*_*A*_ male for *F. cinerea* in this comparison. We used these individuals to visualize mapping quality along chromosome 3. To do so, we converted the BAM outputs from bwa-mem2 to SAM format with “view” in SAMtools version 1.18 (Li et al. 2009) and extracted the mapping quality for each mapped position. We then used a custom script to calculate the mean mapping quality in 10kbp windows in R version 4.2.3 (R Core Team 2023), and visualized the results with ggplot2 (Wilkinson 2011). We identified windows with mapping quality < 40 (*n* = 318, Figure S5) and omitted windowed phylogenies that fell into these poor mapping regions. After omitting windows in poor mapping regions, we retained 6,605 (88.40%) of the initially identified 7,472 50-SNP tree windows.

## Results

### *An* M_D_ *haplotype is present in the* pactrans *clade, the* glacpodz *clade, and* F. cinerea

In allopatric populations of *F. pacifica, F. transmontanis*, and *F*. cf. *transmontanis*, we identified two clusters that contained most individuals, putatively *M*_*D*_*/M*_*A*_ (*n* = 10, 18, 10) and *M*_*A*_*/M*_*A*_ (*n* = 31, 58, 15), respectively (Figure 2A-C). We identified one *F. pacifica* worker with a putative *M*_*D*_*/M*_*D*_ genotype (Figure 2A). In *F. glacialis*, we identified three haplotypes present in the following combinations, *M*_*D*_/*M*_*A*_ (*n* = 12), *M*_*A*_*/M*_*A*_ (*n* = 25), *P*_*S*_/*M*_*A*_ (*n* = 10), and one individual with the *P*_*S*_/*M*_*D*_ genotype (Figure 2D). We also sampled three *F. glacialis* males with the *M*_*A*_ haplotype (Figure 2D). We used a subset of samples with known supergene genotypes from Purcell et al. 2025 for *F. podzolica* and confirmed their identities as *M*_*A*_/*M*_*A*_ (*n* = 66), *M*_*D*_/*M*_*A*_ (*n* = 31), and *P*_*L*_/*M*_*A*_ (*n* = 36) (Figure 2E). We resequenced *F. cinerea* haploid males and diploid workers with known supergene genotypes (Scarparo et al. 2023). We confirmed the identity of the supergene haplotypes: males were *M*_*A*_ (*n* = 11), *P*_*1*_ (*n* = 10), or *P*_*2*_ (*n* = 11), and the workers (*n* = 13) were *M*_*D*_*/M*_*A*_ (Figure 2F). We note that the *M*_*D*_*/M*_*A*_ workers and the *M*_*A*_ males clustered with each other on PC2 but were distinguished on PC3.

### *The* pactrans *clade* M_D_ *haplotype has a similar gyne-producing function as previously described* M_D_ *haplotypes*

Previous work established that an *M*_*D*_ haplotype was associated with colony gyne production in two Nearctic species, *F. glacialis* and *F. podzolica* (Lagunas-Robles et al. 2021). Here, we found an *M*_*D*_ haplotype in three additional Nearctic species, *F. pacifica, F. transmontanis*, and *F*. cf. *transmontanis*. Despite the shared *M*_*D*_ haplotypes between the *glacpodz* clade and the *pactrans* clade, there were slight differences in the association between genotype and sex ratio phenotype. In the *glacpodz* clade, colonies with the *M*_*D*_/*M*_*A*_ genotype produced gynes, while colonies with the *M*_*A*_/*M*_*A*_ genotype produced males (Lagunas-Robles et al. 2021). In the *pactrans* clade, we found a significant association between colonies with the putative *M*_*D*_ haplotype and gyne production (p = 3.39×10^-5^). However, colonies with the heterozygous *M*_*D*_/*M*_*A*_ genotype produced both gynes and males, while colonies with only the *M*_*A*_/*M*_*A*_ genotype produced males (Table S3). In the Palearctic species, *F. cinerea*, the presence of the *M*_*D*_ haplotype was significantly associated with colony gyne production in monogyne colonies (Scarparo et al. 2023).

### *The Nearctic* M_D_ *haplotypes have distinct boundaries from the* F. cinerea M_D_ *haplotype*

We found that the *F. pacifica, F. transmontanis*, and *F*. cf. *transmontanis M*_*D*_ haplotypes shared similar boundaries to the *M*_*D*_ haplotypes found in the *glacpodz* clade (2-7.5Mbp, Figure 2G-I). Notably, we did not observe a *P* haplotype in any individuals sequenced from the *pactrans* clade (Fig S3). We found that the *F. cinerea M*_*D*_ haplotype was reduced in length compared to congeneric *M*_*D*_ haplotypes with approximate boundaries at 2.67Mbp and 7.5Mbp on chromosome 3 (Figure 2L). The right-hand boundary occurs at approximately 7.5Mbp, but the abundance of repetitive sequences in that region resulted in low mapping quality, making it difficult to pinpoint the boundary (Figure S4). We observed no differentiated regions in other parts of the genome that were associated with supergene variation on chromosome 3, aside from the chromosome 9 supergene previously described in *F. cinerea* (Scarparo et al. 2023).

### *The topological history of the* M_D_ *haplotypes suggests that the Nearctic* M_D_ *haplotypes and the* F. cinerea M_D_ *haplotype have different origins*

We examined the evolutionary history of the *M*_*D*_ haplotype by assigning each 50-SNP window to a criterion topology. A large number of windows in the 2-7.5Mbp region of chromosome 3 were shared between the *M*_*D*_ haplotypes of all five Nearctic species, and very few windows were shared between the *M*_*D*_ haplotypes of *F. cinerea* and the Nearctic species (Figure 3; Table 1). This suggests that the Nearctic *M*_*D*_ haplotypes and the shorter *F. cinerea M*_*D*_ haplotype have very little shared genetic variation, consistent with the distinct boundaries identified by the putative inversion breakpoint.

**Table 1.**
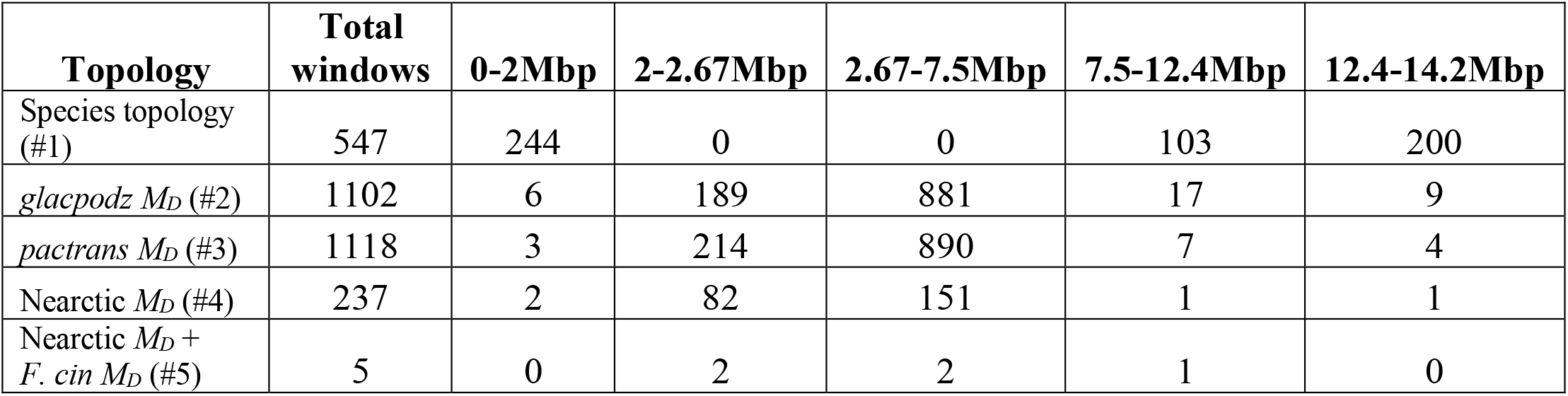
The count of 50-SNP windows for each criteria clade topology in Figure 1. The boundaries for the Nearctic *M*_*D*_ haplotypes are 2 and 7.5Mbp and the shorter *F. cinerea M*_*D*_ haplotype is between 2.67Mbp and 7.5Mbp. In the Nearctic species, the *M*_*A*_ haplotype and the *M*_*D*_ haplotype recombine between 0-2Mbp and 7.5-14.2Mbp. In *F. cinerea*, the *M*_*A*_ and the *M*_*D*_ haplotype also recombine between 2-2.67Mbp.

**Figure 3.**
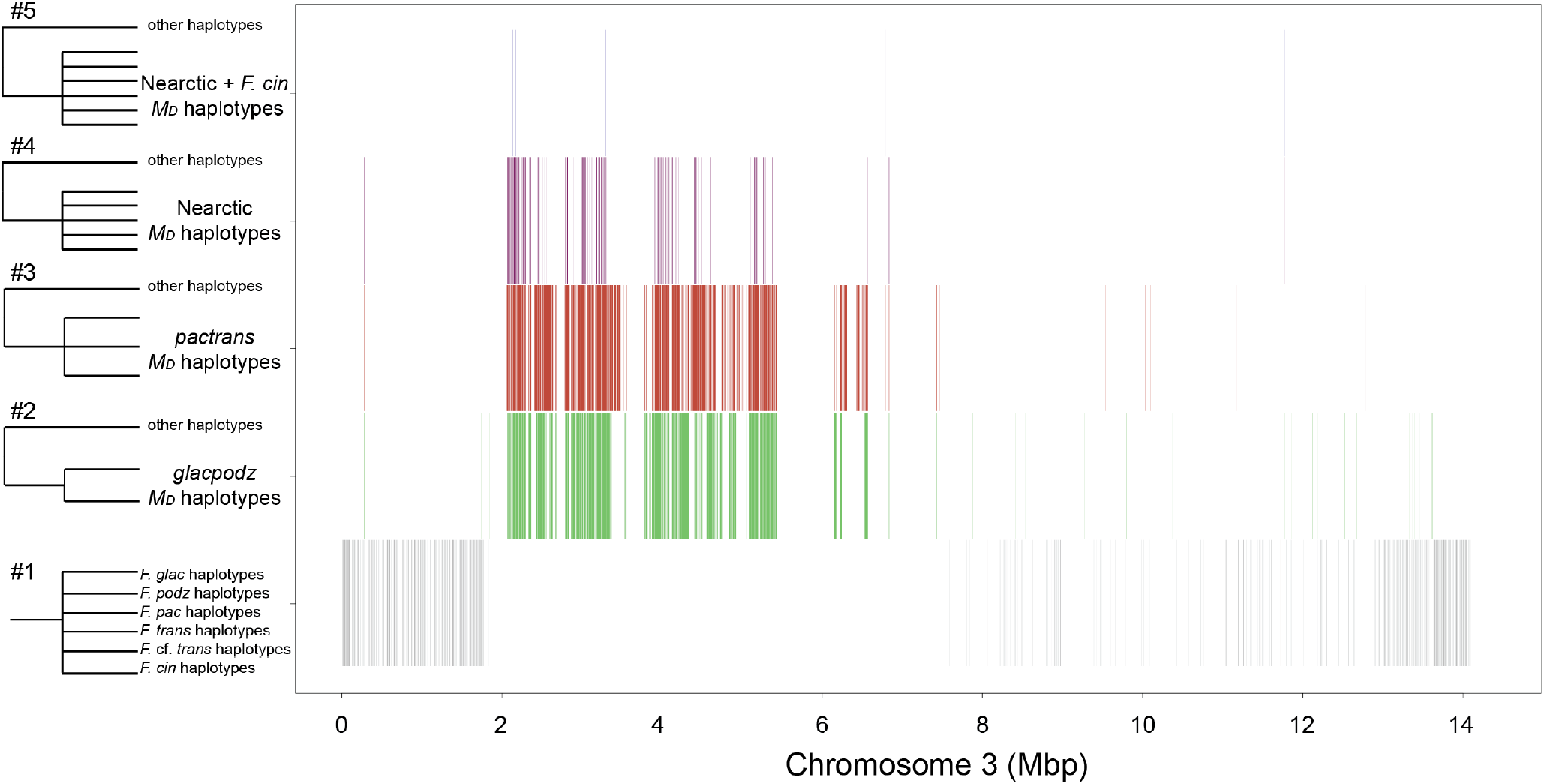
Ahaplotype topological analysis of 50-SNP windows along chromosome 3 suggests that the Nearctic *M*_*D*_ haplotypes are more similar to each other than to the *F. cinerea M*_*D*_ haplotype. Criteria clades, represented by the polytomies illustrated in the left column, were searched for among the haplotype topologies on chromosome 3. When the windowed topology matched a criteria clade, the window was plotted based on its physical position. For topologies matching clade #1, individuals from each species form a clade regardless of their supergene haplotype. This pattern was found outside of the *M*_*D*_ haplotype region (2-7.5Mbp). The remaining criteria clades represented polytomies with specific *M*_*D*_ haplotype relationships: the *glacpodz* clade *M*_*D*_ haplotypes (#2), the *pactrans* clade *M*_*D*_ haplotypes (#3), the Nearctic *M*_*D*_ haplotypes (#4), and all six *M*_*D*_ haplotypes (#5). Many windows within the 2-7.5Mbp region clustered Nearctic *M*_*D*_ haplotypes (#2-4), while very few windows clustered both Nearctic and Palearctic *M*_*D*_ haplotypes (#5). The number of windows matching each criteria clade is reported in Table 1.

### *The role of the* P *haplotype in the formation of multiple* M_D_ *haplotypes*

To test whether the *M*_*D*_ haplotype originated from a *P* haplotype, we leveraged the *F. cinerea P*_*1*_ and *P*_*2*_ haplotypes and the *F. podzolica P*_*L*_ haplotype. We found more windows within the 2.67-7.5Mbp region that formed a clade between the *F. cinerea M*_*D*_ haplotype and the *F. cinerea P*_*1*_ haplotype (*n* = 380) than the *F. cinerea M*_*D*_ and *F. cinerea P*_*2*_ haplotype (*n* = 76) suggesting that the *M*_*D*_ haplotype in *F. cinerea* originated from the *P*_*1*_ haplotype after the divergence between the *P*_*1*_ haplotype and *P*_*2*_ haplotype (Figure 4, Table 2). Searching for clades containing the *F. podzolica P*_*L*_ haplotype and the Nearctic *M*_*D*_ haplotypes, we recovered 108 windows within the 2-7.5Mbp region (Figure 4). We detected far fewer windows with topologies grouping the *F. podzolica P*_*L*_ with the *F. cinerea M*_*D*_ or the Nearctic *M*_*D*_ with the *F. cinerea P*_*1*_ and *P*_*2*_ (Figure 4, Table 2), and most of these windows clustered outside of the *M*_*D*_ region in both comparisons. Note that 11 of 15 windows grouping the Nearctic *M*_*D*_ with the *F. cinerea P*_*1*_ and *P*_*2*_ occurred between 2-2.67Mbp, which is outside of the *F. cinerea M*_*D*_ haplotype boundary. Taken together, these results suggest that the *F. cinerea M*_*D*_ haplotype and the Nearctic *M*_*D*_ haplotypes originated from different *P* haplotypes.

**Table 2.** The count of 50-SNP windows for each criteria clade topology in Figure 4. The boundaries for the Nearctic *M*_*D*_ haplotypes are from 2-7.5Mbp and the reduced *F. cinerea M*_*D*_ haplotype has boundaries from 2.67-7.5Mbp. All the *P* haplotypes in this analysis share boundaries between 2-12.4Mbp. The regions with free recombination between all haplotypes are located between 0-2Mbp and 12.4-14.2Mbp.

| Topology | Total windows | 0-2Mbp | 2-2.67Mbp | 2.67-7.5Mbp | 7.5-12.4Mbp | 12.4-14.2Mbp |
| --- | --- | --- | --- | --- | --- | --- |
| *F. cin* $P_1$ + *F. cin* $M_D$ (#6) | 636 | 23 | 18 | 380 | 176 | 39 |
| *F. cin* $P_2$ + *F. cin* $M_D$ (#7) | 257 | 30 | 8 | 76 | 98 | 45 |
| *F. cin* $P_1$, $P_2$ + Nearctic $M_D$ (#8) | 15 | 0 | 11 | 4 | 0 | 0 |
| *F. podz* $P_L$ + Nearctic $M_D$ (#9) | 109 | 0 | 43 | 65 | 0 | 1 |
| *F. podz* $P_L$ + *F. cin* $M_D$ (#10) | 41 | 0 | 0 | 6 | 33 | 2 |
| *F. cin* $P_1$, $P_2$ + *F. podz* $P_L$ (#11) | 478 | 1 | 11 | 65 | 386 | 15 |

**Figure 4.**
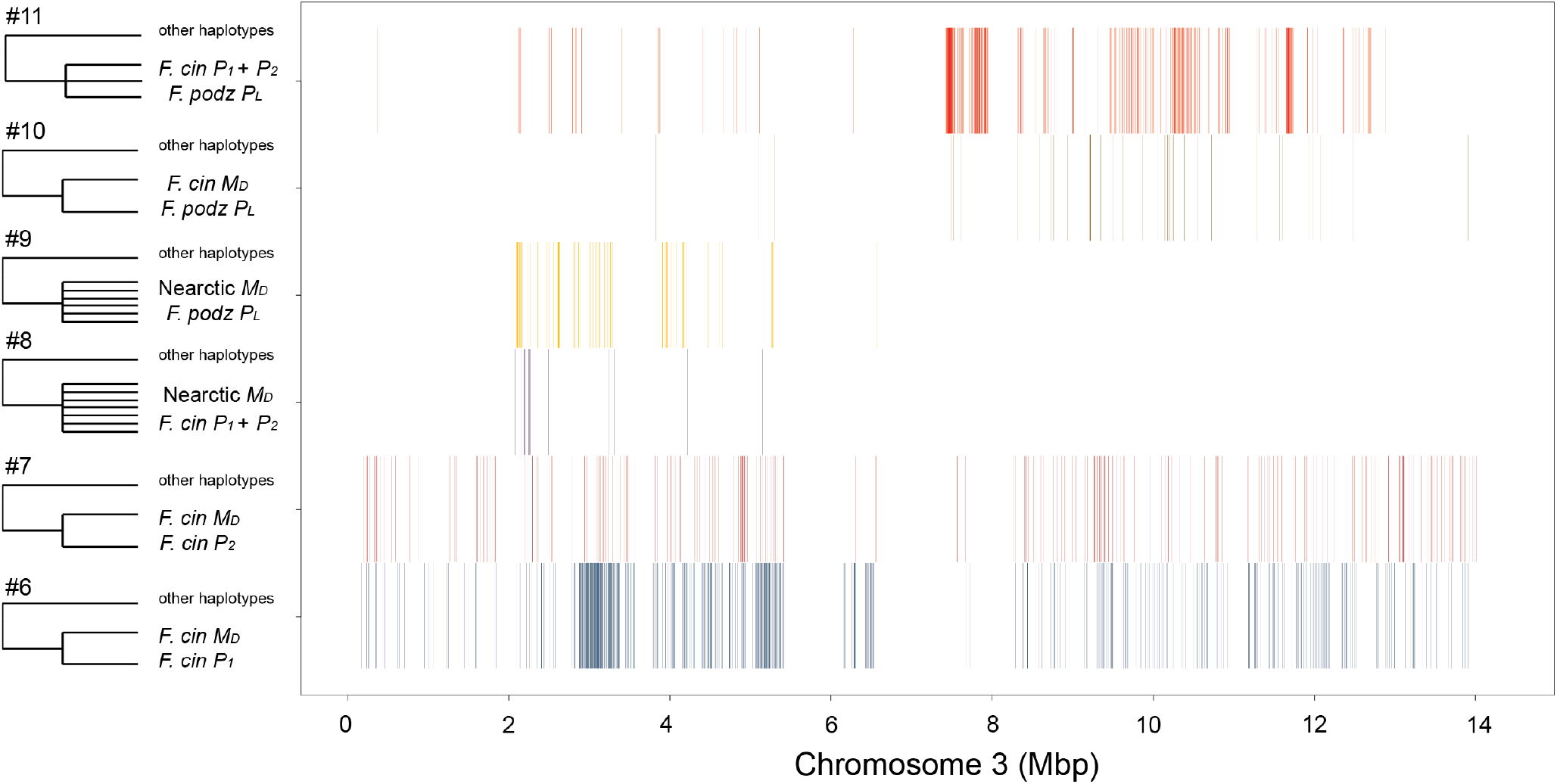
Atopological analysis of 50-SNP windows meeting criteria clades that inform the relationships between *M*_*D*_ haplotypes and the *P* haplotypes. Criteria clades, represented by polytomies in the left column (#6-11), were searched for among the haplotype topologies on chromosome 3. When the windowed topology matched a criteria clade, the window was plotted based on its physical position. There were more windows wherein the Nearctic *M*_*D*_ haplotypes grouped with the *F. podzolica P*_*L*_ haplotype (#9) than with the *F. cinerea P*_1_ and *P*_2_ haplotypes (#8). Along the same lines, there were more windows wherein the *F. cinerea M*_*D*_ haplotype grouped with the *F. cinerea P*_1_ (#6) than the *F. cinerea P*_2_ (#7). Furthermore, there were only six windows wherein the *F. cinerea M*_*D*_ haplotype grouped with the *F. podzolica P*_*L*_ haplotype (#10). These patterns suggest that the Nearctic *M*_*D*_ haplotype and the *F. cinerea M*_*D*_ haplotype have independent origins from different *P* haplotypes. The *F. podzolica* and *F. cinerea P* haplotypes grouped together in windows spanning the 2-12.4Mbp supergene region, with a higher concentration of windows in the 7.5-12.4Mbp section of the supergene (#11). The number of windows matching each criteria clade is reported in Table 2.

Lastly, we searched for SNP trees in which all the *P* haplotypes formed a clade (Figure 4, Table 2). We found 96.65% of the windows containing the *P* haplotype clades between 2-12.4Mbp (Figure 4, Table 2), the region that typically differentiates the *M* and *P* haplotypes in other *Formica* species (Brelsford et al. 2020). We did not find evidence suggesting the *M*_*D*_ haplotypes originated from a rearrangement of the ancestral *M* haplotype (Figure S6).

## Discussion

### *On the origins of the* M_D_ *haplotype and its link to an ancient queen number supergene*

We find evidence for two origins of the *M*_*D*_ haplotype – one origin in the Nearctic *Formica* species in this study (Figure 3) and another in *F. cinerea* or an ancestor of this Palearctic species, hereafter called “the *F. cinerea* lineage” (Figure 4). All Nearctic species share boundaries of the *M*_*D*_ haplotype on chromosome 3 at approximately 2-7.5Mbp (Figure 2G-K). In contrast, *F. cinerea* has *M*_*D*_ haplotype boundaries at approximately 2.67-7.5Mbp (Figure 2L). Additionally, the *M*_*D*_ haplotypes of all five Nearctic species cluster together in a substantial number of windows (*n* = 233, Table 1, Figure 3) within the *M*_*D*_ haplotype boundaries. Within the boundaries of the *F. cinerea M*_*D*_ haplotype, however, only two windows grouped the *M*_*D*_ of all six species in the study (Figure 3, Table 1).

Both origins of the *M*_*D*_ haplotypes were apparently derived from a *P* haplotype. The Nearctic *M*_*D*_ haplotypes group with the *F. podzolica P* haplotype over much of their length (Figure 4, Table 2). Similarly, the *F. cinerea M*_*D*_ haplotype grouped with one of the two *F. cinerea P* haplotypes, with a concentration of shared topology windows in the 2.67-7.5Mbp region of chromosome 3 (Figure 4, Table 2).

We infer that the Nearctic *M*_*D*_ haplotype formed in an ancestor of the *pactrans* and *glacpodz* clades, since the *M*_*D*_ haplotypes of all five species have shared boundaries, group together in topologies across the extent of the *M*_*D*_ haplotype, and all share topologies with the *F. podzolica P*_*L*_ in the 2-7.5Mbp region of chromosome 3. The *F. cinerea M*_*D*_ haplotype formed more recently than the split between the *F. cinerea P*_*1*_ and *P*_*2*_ haplotypes, since it clustered with the *P*_*1*_ in many more windows than with the *P*_*2*_. The difference in the extent of the *M*_*D*_ haplotypes in the Nearctic species and *F. cinerea* and the shared function of the supergene haplotypes suggest that genes in the 2-2.67Mbp region likely do not control gyne production.

One limitation of our study is having only one Palearctic species in our analyses. It is possible that the version of the *M*_*D*_ haplotype found in the *F. cinerea* lineage is present in other Palearctic relatives. Sampling additional Palearctic species will be necessary to unravel the history of the Palearctic *M*_*D*_ haplotype and understand how it relates to a gyne-producing function.

### A *convergent recombinant origin of the* M_D_ *haplotype*

Recombination events between pre-existing supergene haplotypes can generate additional haplotypes (De-Kayne et al. 2025). In the African monarch butterfly, *Danaus chrysippus*, color pattern is determined by six divergent haplotypes (De-Kayne et al. 2025). Ancestry and admixture patterns near or at the promoter region of two genes suggest that one of the haplotypes is a recombinant product of two pre-existing haplotypes (De-Kayne et al. 2025). The recombinant origin of the African monarch butterfly could parallel the *Formica M*_*D*_ haplotype, where a portion of a *P* haplotype likely recombined onto an *M* haplotype background (Figure 4). This led to two diverging *M* haplotypes, the proto-*M*_*D*_ haplotype and the *M*_*A*_ haplotype.

In ants, rare recombination events between supergene haplotypes likely contribute to phenotypic novelty in multiple systems. Interestingly, recombination between the *M* and the *P* haplotypes likely led to the movement of two *P*-specific genes onto an *M* haplotype background in *F. aquilonia* and *F. paralugubris* (Sigeman et al. 2024). Sigeman et al. suggested that strong selection against the *P* haplotype could have favored the fixation of this recombinant *M* haplotype. The near absence of the *P*/*M*_*D*_ genotype is notable in all species with an *M*_*D*_ haplotype (Lagunas-Robles et al. 2021, Scarparo et al. 2023, Purcell et al. 2025, this study). We speculate that the recombinant origin of the *M*_*D*_ haplotype resulted in some of the *P* haplotype genetic load being moved onto the *M*_*D*_ haplotype, resulting in a deleterious *P*/*M*_*D*_ genotype combination. Additional sampling of polygyne colonies with split sex ratios is needed to unravel the causes of the paucity of the *P/M*_*D*_ haplotype.

*The loss of a queen number supergene in* F. pacifica, F. transmontanis, *and* F. *cf*. transmontanis Genetic work on colony queen number in the *Formica* genus has shown that variation in colony social form is typically maintained by *M* and *P* haplotypes. Colonies are polygyne when workers have at least one copy of a *P* haplotype (Purcell et al. 2014, Brelsford et al. 2020, Lagunas-Robles et al. 2021, McGuire et al. 2022, Pierce et al. 2022, Scarparo et al. 2023 but see Sigeman et al. 2024 and Lagunas-Robles et al. 2025). In the species *F. pacifica, F. transmontanis*, and *F*. cf. *transmontanis*, we do not find any individuals with a *P* haplotype. If all three species are indeed lacking a *P* haplotype and are confirmed to be strictly monogyne with a robust within-colony sampling approach, this *Formica* clade has reverted from socially polymorphic to strictly monogyne and also lost the *P* haplotype. Whether the species in the *pactrans* clade are strictly monogyne is an open question.

### Avenues for future sex ratio studies

Based on the initial discovery of the sex ratio supergene in *Formica* (Lagunas-Robles et al. 2021), we expected that monogyne colonies headed by queens with the *M*_*D*_ haplotype would produce gynes and colonies with *M*_*A*_*/M*_*A*_ queens would produce males. However, in the *pactrans* clade, a species where we only identified *M* haplotypes, the *M*_*D*_ haplotype was found in colonies producing both gynes and males, while the *M*_*A*_ haplotype was homozygous in male-producing colonies. Lagunas-Robles et al. proposed a verbal model in *F. glacialis* in which a heterozygous *M*_*D*_/*M*_*A*_ queen mates with an *M*_*A*_ haploid male leading to *M*_*D*_/*M*_*A*_ or *M*_*A*_/*M*_*A*_ queens. This simple mechanism would lead to gyne-producing colonies with a *M*_*D*_/*M*_*A*_ queen and male-producing colonies with a *M*_*A*_/*M*_*A*_ queen. This model would not require recessive lethality of the *M*_*D*_ haplotype to explain the lack of *M*_*D*_/*M*_*D*_ genotypes in the population as *M*_*D*_ males should be rare. In contrast, the *M*_*D*_ haplotype in the *pactrans* clade is present in colonies producing both gynes and males. This observation suggests that *M*_*D*_ males should be produced at a higher rate than in *F. glacialis*, and that these males should produce some *M*_*D*_/*M*_*D*_ worker offspring when mated with heterozygous queens. However, we only observed a single *M*_*D*_/*M*_*D*_ worker in our *pactrans* samples (Figure 2A-C). This paucity of *M*_*D*_*/M*_*D*_ homozygotes is consistent with the low frequency of *P/M*_*D*_ genotypes in other species, and is aligned with our speculation that some of the genetic load from an ancestral *P* haplotype may still be present on the *M*_*D*_. Functionally, the differences among species in the sex ratio of offspring produced by *M*_*D*_*/M*_*A*_ queens could point to different mechanisms maintaining colony sex ratio in the *pactrans* clade and *glacpodz* clade despite sharing a Nearctic *M*_*D*_ haplotype.

## Conclusion

The evolutionary processes that lead to the diversification of autosomal supergenes can be difficult to disentangle as supergene haplotypes are often limited to a single species. Examining convergent polymorphisms can elucidate the evolutionary processes that lead to supergenes. For example, in anther-smut fungi, a “young” (0.5-3.2 million years) mating-type locus supergene expanded through convergent evolution of suppressed recombination at least nine times (Branco et al. 2018; Duhamel et al. 2022). Additionally, the presence of many haplotypes can help uncover how supergene systems diversify. In the African monarch butterfly, one of the six described haplotypes underlying color pattern is likely a recombinant haplotype (De-Kayne et al. 2025). Here, we find additional support for the formation of new haplotypes resulting from recombination between pre-existing supergene haplotypes. We also show that a recombinant haplotype can shape a distinct function compared to its pre-existing counterparts. The recombination events between the ancient *M* and *P* haplotypes (~23 million years old, Purcell et al. 2021) resulted in a recombinant haplotype, the *M*_*D*_, that is associated with colony-level gyne production. Strikingly, we find that convergent *M*_*D*_ haplotypes likely arose twice from different *P* haplotypes, once in the ancestor of the Nearctic *Formica* species featured in this study and once in the *F. cinerea* lineage.

## Acknowledgements

We thank Polly Campbell, and David Reznick for comments on an earlier version of this manuscript. Giulia Scarparo and Marie Palanchon graciously shared DNA samples for *F. cinerea*. Zul Alam helped to generate the sequence data. Felicia Lin, Marie Palanchon, Daniel Pierce, and Elisa Seals helped collect *F. pacifica, F*. cf. *transmontanis*, and *F. transmontanis* samples. Darin McGuire and Junxia Zhang helped to collect *F. glacialis* samples. Madison Sankovitz performed the DNA extractions for *F. glacialis* samples for this study. This material is based upon work supported by the United States National Science Foundation Graduate Research Fellowship to G.L.-R. under Grant No. DGE-1326120, US National Science Foundation CAREER to J.P under DEB Grant No. 1942252, and DEB Grant No. 1754834 to A.B. and J.P. G.L.-R. received funding from the Vaughan H. Shoemaker Graduate Fund at University of California, Riverside to conduct field work. Computations were performed using the computer clusters and data storage resources of the High-Performance Computing Center at University of California, Riverside, which were funded by grants from United States National Science Foundation (MRI-2215705, MRI-1429826) and United States National Institute of Health (1S10OD016290-01A1).

## Data Availability

Sequence data will be made available on NCBI upon acceptance. Data and R scripts used to generate plots will also be made available upon acceptance.

## Notes

### Competing Interest Statement

The authors have declared no competing interest.

