## Supplemental Materials for "Multiple origins of a sex ratio supergene in *Formica ants*"

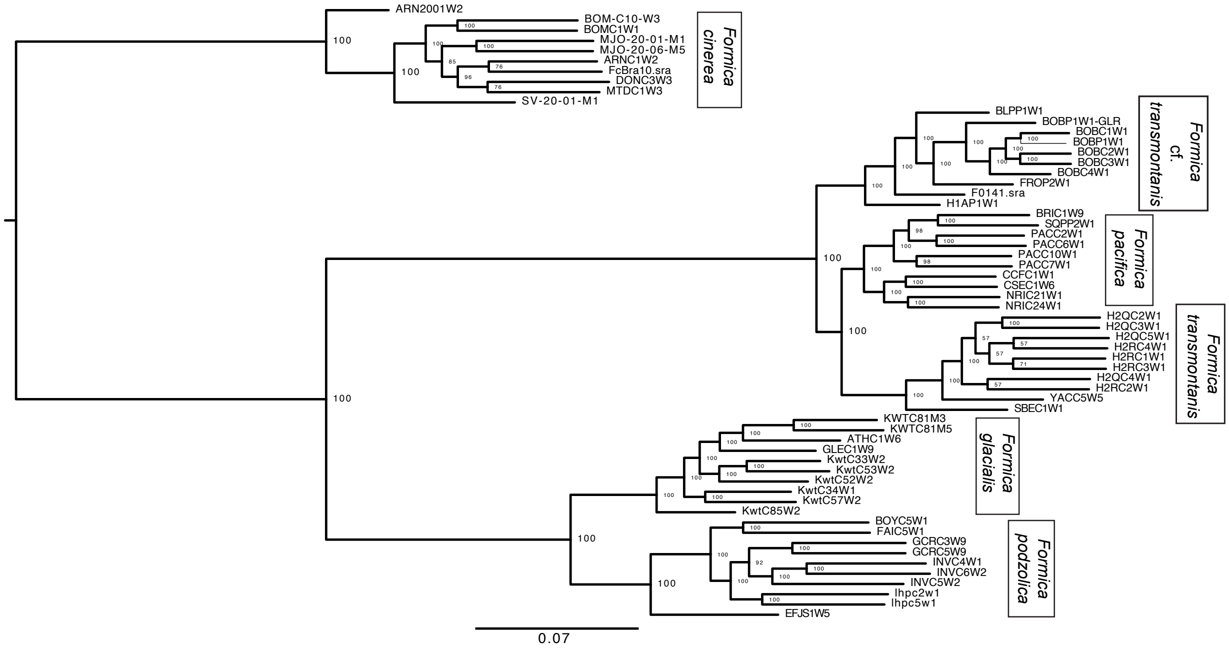


**Figure S1.** We used a subset of 10 high coverage genomes (mean depth = 5.76 - 24.69) for each putative species group to generate a maximum-likelihood tree. We excluded chromosome 3 in this analysis due to its trans-species supergene polymorphism. Species splits had 100% bootstrap support. Of note, we found that *F. transmontanis* as currently described is paraphyletic: a northern lineage we are delineating as *F. transmontanis* is sister to *F. pacifica*, and a southern lineage we are delineating as *F.* cf. *transmontanis* diverged prior to the common ancestor of *F. transmontanis* and *F. pacifica*.

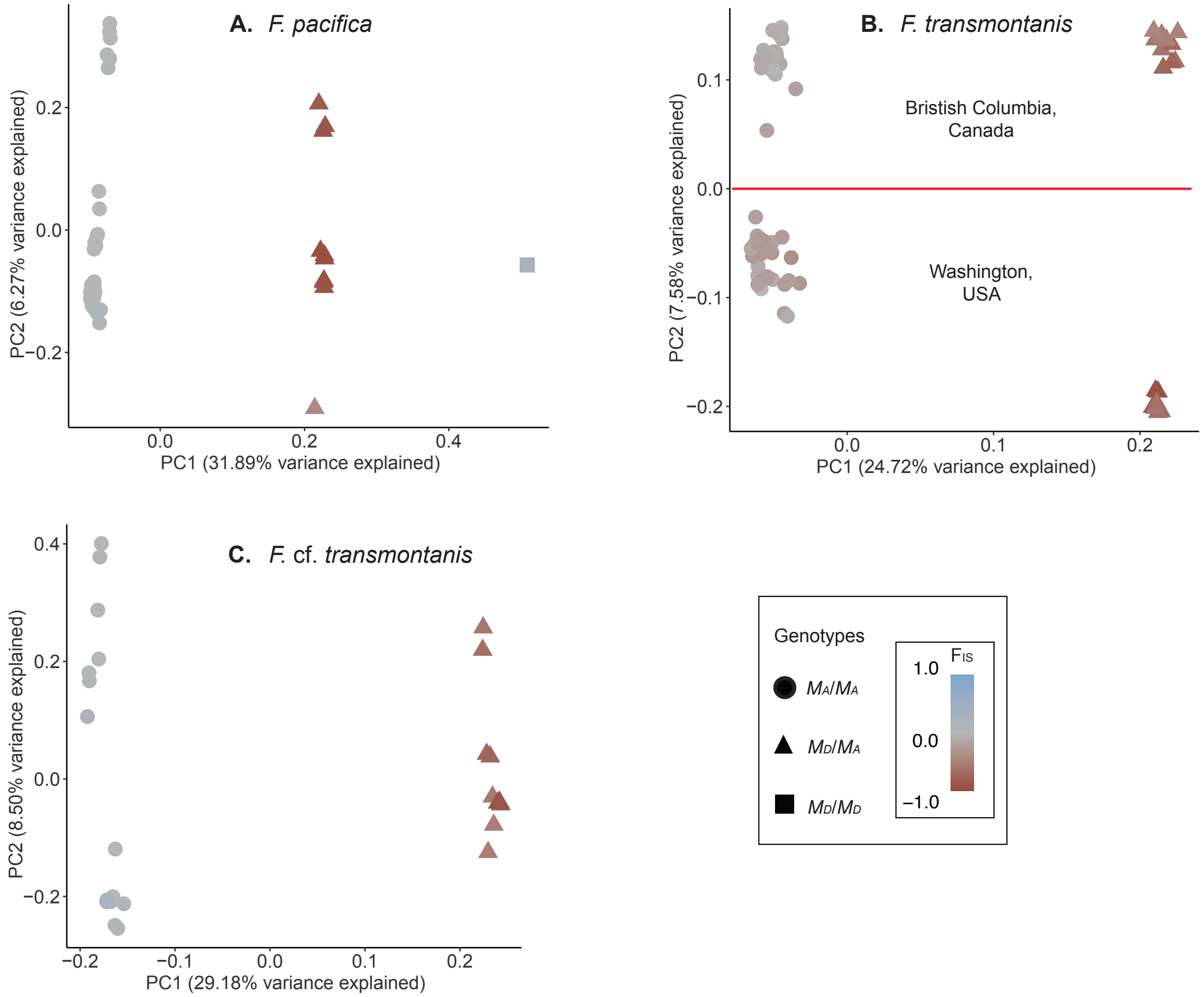


**Figure S2.** Principal component analyses were performed on SNPs found on chromosome 3 between 2-7.5Mbp. Heterozygosity ($F_{IS}$) was estimated for the same region for *F. pacifica*, *F. transmontanis*, and *F.* cf. *transmontanis*. A combination of $F_{IS}$ and PC1 was used to determine the genotypes assigned in Figure 1 in the main text. **(A)** *F. pacifica* has three genetic clusters separating on PC1. This pattern is expected when supergenes are present in the heterozygous and both homozygous states. The middle cluster had negative $F_{IS}$ values suggesting these individuals were heterozygous for the 2-7.5Mbp supergene region. **(B)** *F. transmontanis* and **(C)** *F.* cf. *transmontanis* both have two clusters along PC1, separating individuals with and without the $M_D$ haplotype. In *F. transmontanis* (**B**), the individuals above the red line were collected in British Columbia, Canada while individuals below the red line were collected from Washington, USA. The negative $F_{IS}$ clusters (right) had the $M_D$ haplotype individuals. In *F.* cf. *transmontanis* (**C**), the cluster with the negative $F_{IS}$ (right) had the $M_D$ haplotype individuals.

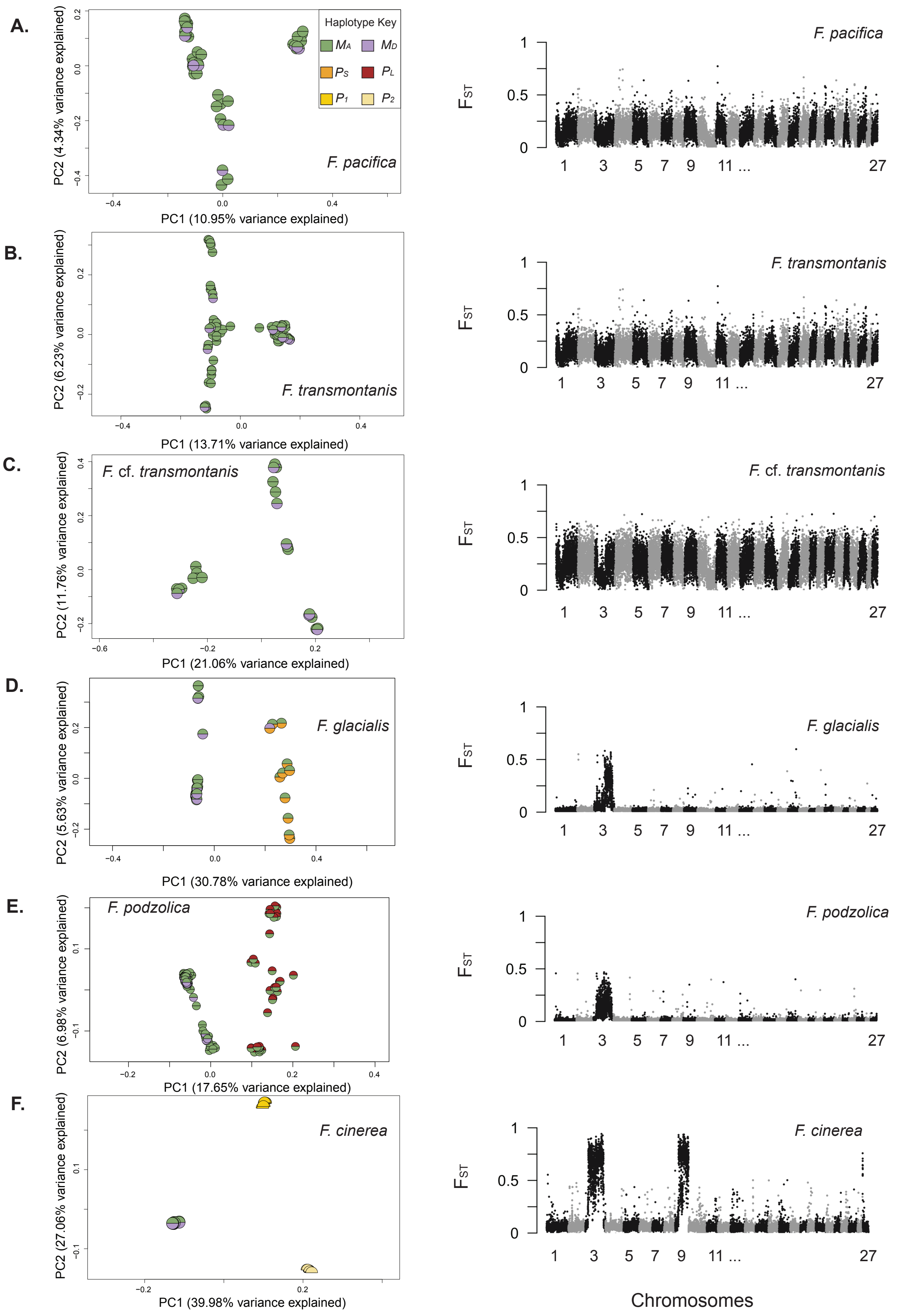
A.
B.
C.
D.
E.
F.
Haplotype Key
MA MD
PS PL
P1 P2
F. pacifica
F. transmontanis
F. cf. transmontanis
F. glacialis
F. podzolica
F. cinerea
PC1 (10.95% variance explained)
PC2 (4.34% variance explained)
PC1 (13.71% variance explained)
PC2 (6.23% variance explained)
PC1 (21.06% variance explained)
PC2 (11.76% variance explained)
PC1 (30.78% variance explained)
PC2 (5.63% variance explained)
PC1 (17.65% variance explained)
PC2 (6.98% variance explained)
PC1 (39.98% variance explained)
PC2 (27.06% variance explained)
FST
1 3 5 7 9 11 ... 27
Chromosomes

**Figure S3. (A-C)** Based on principal component analyses of chromosome 3 between 7.5Mbp and 12.4Mbp, we found no evidence of a *P* haplotype in *F. pacifica*, *F. transmontanis*, or *F.* cf. *transmontanis*. When we estimated $F_{ST}$ between the most distant clusters along PC1, we recovered noisy patterns that suggested the SNPs found in the 7.5-12.4Mbp region were not informative of supergene genotypes. **(D-F)** We recovered known *P* haplotype variation in *F. glacialis* (Lagunas-Robles et al. 2021), *F. podzolica* (Purcell et al. 2025), and *F. cinerea* (Scarparo et al. 2023).

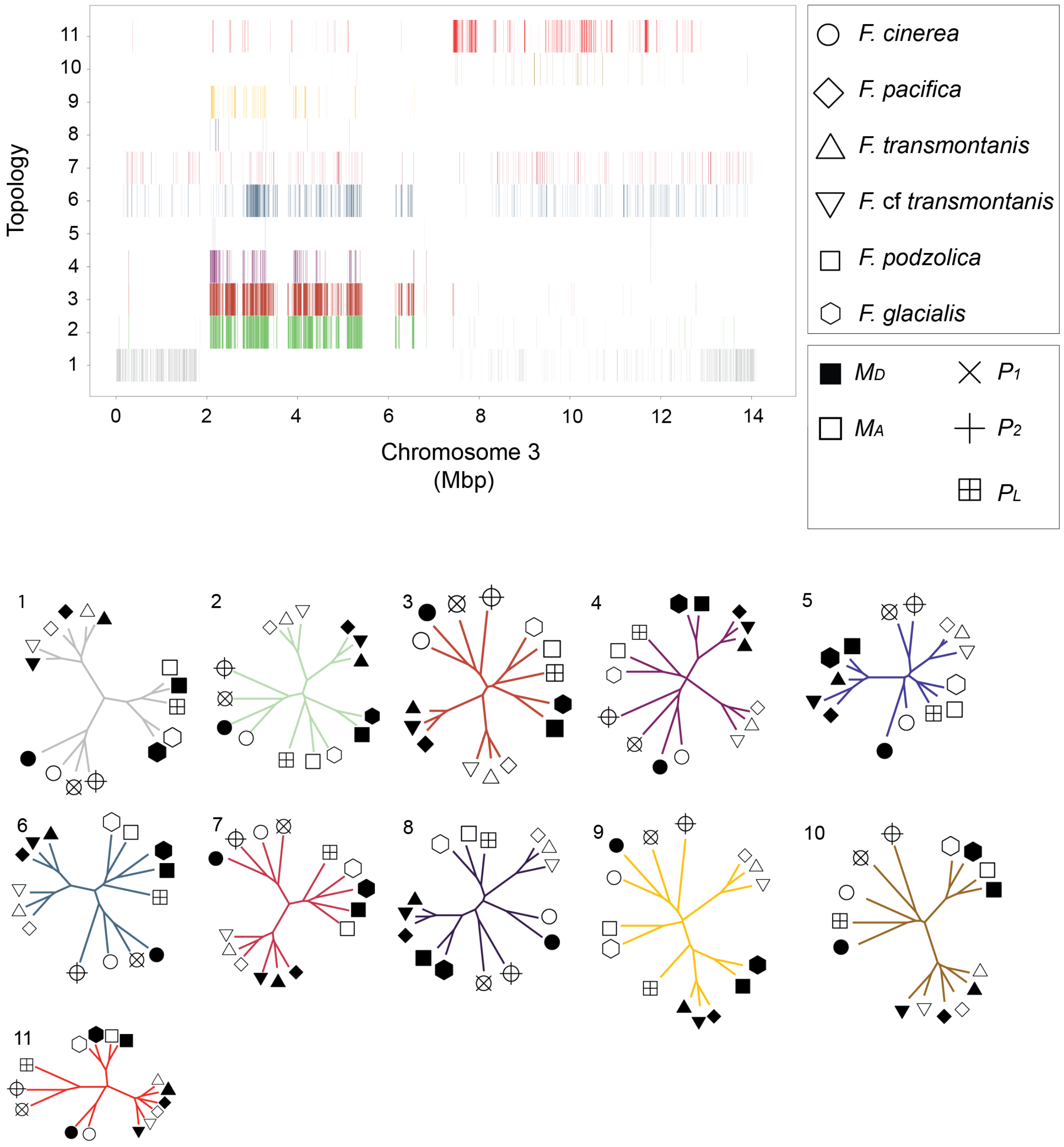


**Figure S4.** In the top panel, each row has 50-SNP windows that meet the specific criteria topology (#1-11), also shown in Figures 3 and 4 of the main text. We extracted the allele frequencies of all SNPs that were mapped for each condition and summarized all the windows in an unrooted neighbor-joining tree (bottom panel). Row numbers match their numbered topology. The shapes represent each species and shape attributes distinguish the haplotypes.

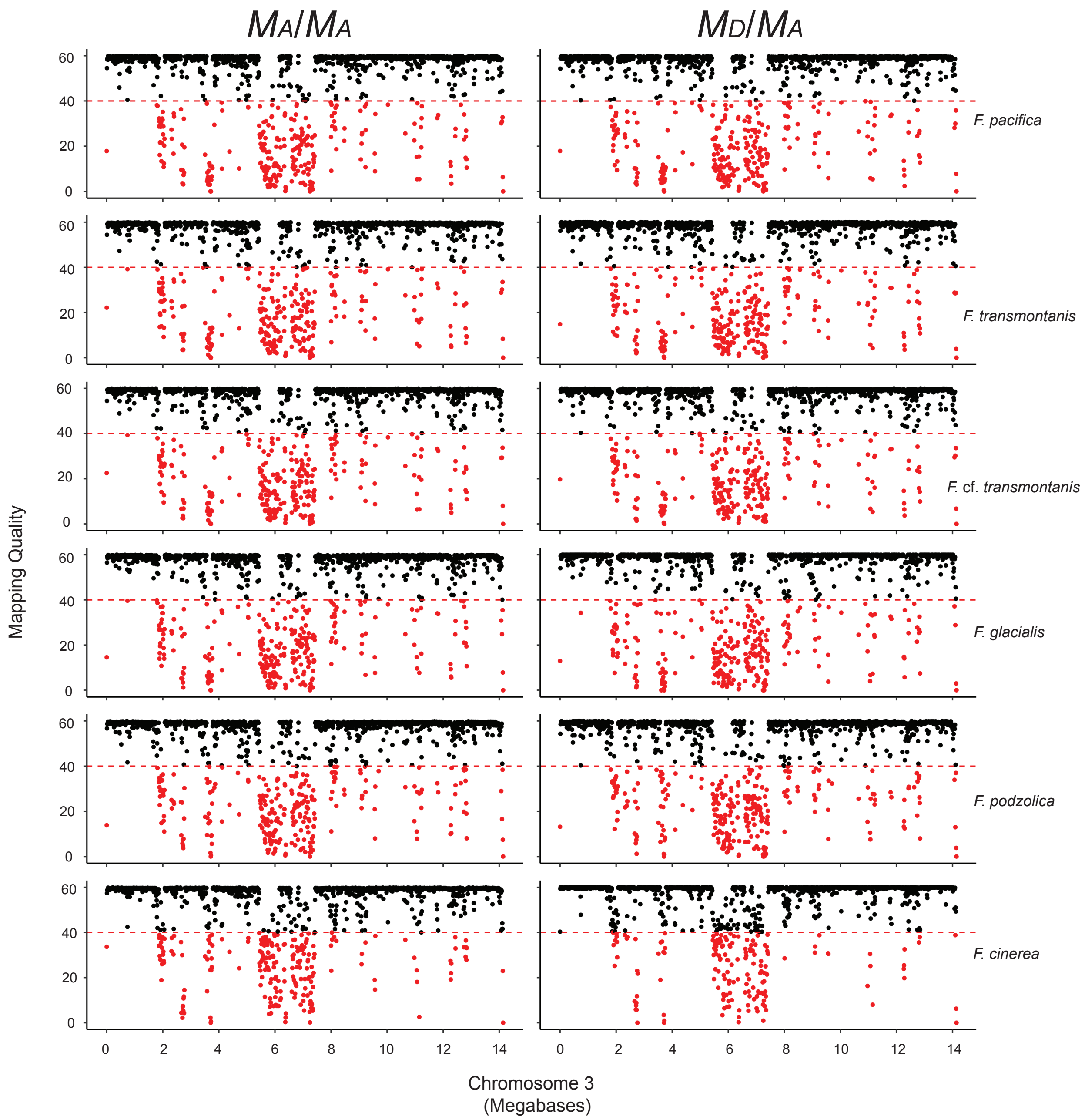


**Figure S5.** We assessed the raw read mapping quality across chromosome 3 in 10kbp windows. We selected two individuals for each species with the highest depth for the $M_A/M_A$ and $M_D/M_A$ genotypes. We selected the highest depth $M_A$ male for *F. cinerea* in this comparison. We found a decrease in read mapping quality from approximately 5.5-6.5Mbp and 7-7.5Mbp across all individuals and genotypes. This suggested that a highly repetitive region interfered with mapping short reads independent of supergene genotype. We estimated the right-hand supergene boundary to be approximately 7.5Mbp, but long-read data would be necessary to pinpoint the exact boundary and whether the $M_D$ haplotypes captured parts of a centromere. We denoted regions of low mapping quality (in red) and omitted these 10-kbp windows from the topology analyses.

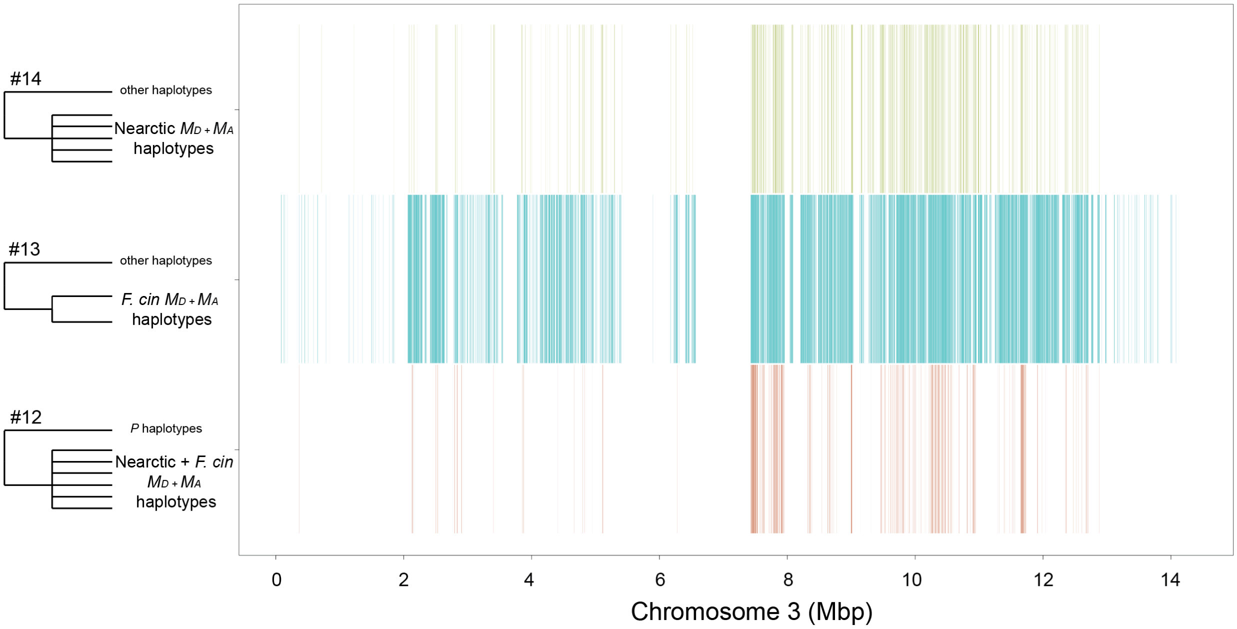


**Figure S6.** A topological analysis of 50-SNP windows meeting criteria clades that inform the relationships between $M_D$ haplotypes and the $M_A$ haplotypes. A high number of topologies are concentrated in the 7.5-12.4Mbp interval (#12-14). This is expected as the ancestral *M* haplotype is present between 7.5-12.4Mbp and is not differentiated between the $M_D$ and $M_A$ haplotypes (Figure 2). The 2-7.5Mbp range has less windows when examining topologies #12 and #14 as these include multiple $M_D$ haplotypes. Topology #13 is within *F. cinerea* and shows a high density of shared topologies across the 2-12.4Mbp region. Taken together, these results suggest that the $M_D$ haplotypes did not result from an $M_A$ haplotype rearrangement.

**Table S1.** Summary of the sequenced species for this study and samples from published studies.

| Species | New data (*n*) | Public data (*n*) | BioProject sources |
|---|---|---|---|
| *F. glacialis* | 41 | 8<br>2 | Lagunas-Robles et al. 2021 (PRJNA759919)<br>Purcell et al. 2021 (PRJNA750082) |
| *F. podzolica* | - | 123<br>10 | Purcell et al. 2025 (PRJNA1293783, *in review*)<br>Lagunas-Robles et al. 2021 (PRJNA759919) |
| *F. pacifica* | 42 | - | |
| *F. transmontanis* | 76 | - | |
| *F.* cf. *transmontanis* | 24 | 1 | Purcell et al. 2021 (PRJNA750082) |
| *F. cinerea* | 43 | 2 | Brelsford et al. 2020 (PRJNA557080) |

**Table S2.** We used sampled individuals from the *pactrans* complex to design an RFLP assay that distinguished the $M_D$ haplotype from the $M_A$ haplotype. We found fixed differences between the $M_D$ haplotype and the $M_A$ haplotype at position 2789194 on chromosome 3 based on allele frequencies. We then used individuals with a minimum depth of six to validate this diagnostic SNP. The genotype at the position was called in 0/1/2 format where the genotype was homozygous for the reference allele, heterozygous, or homozygous for the alternative allele, respectively. We found one mismatch (in bold) out of 64 samples. This was likely due to a genotyping error. See the Methods for more details.

| Sample ID | Species | Supergene genotype | Scaffold03 2789194 |
|---|---|---|---|
| BRIC1W9 | *F. pacifica* | $M_A/M_A$ | 2 |
| BRIC2W9 | *F. pacifica* | $M_A/M_A$ | 2 |
| BRIC4W9 | *F. pacifica* | $M_A/M_A$ | 2 |
| BRIC7W9 | *F. pacifica* | $M_A/M_A$ | 2 |
| CCFC1W1 | *F. pacifica* | $M_A/M_A$ | 2 |
| CCFC3W1 | *F. pacifica* | $M_D/M_A$ | 1 |
| CSEC1W6 | *F. pacifica* | $M_A/M_A$ | 2 |
| CSEP1W6 | *F. pacifica* | $M_D/M_A$ | 1 |
| HOPC10W1 | *F. pacifica* | $M_A/M_A$ | 2 |
| HOPC11W1 | *F. pacifica* | $M_A/M_A$ | 2 |
| HOPC1W1 | *F. pacifica* | $M_D/M_A$ | 1 |
| HOPC2W1 | *F. pacifica* | $M_A/M_A$ | 2 |
| HOPC3W1 | *F. pacifica* | $M_A/M_A$ | 2 |
| HOPC4W1 | *F. pacifica* | $M_A/M_A$ | 2 |
| HOPC5W1 | *F. pacifica* | $M_A/M_A$ | 2 |
| HOPC6W1 | *F. pacifica* | $M_A/M_A$ | 2 |
| HOPC7W1 | *F. pacifica* | $M_A/M_A$ | 2 |
| HOPC8W1 | *F. pacifica* | $M_A/M_A$ | 2 |
| HOPC9W1 | *F. pacifica* | $M_D/M_A$ | 1 |
| NRIC20W1 | *F. pacifica* | $M_A/M_A$ | 2 |
| **NRIC21W1** | ***F. pacifica*** | $\boldsymbol{M_A/M_A}$ | **1** |
| NRIC22W1 | *F. pacifica* | $M_A/M_A$ | 2 |
| NRIC23W1 | *F. pacifica* | $M_D/M_A$ | 1 |
| NRIC24W1 | *F. pacifica* | $M_A/M_A$ | 2 |
| PACC10W1 | *F. pacifica* | $M_D/M_A$ | 1 |
| PACC1W1 | *F. pacifica* | $M_A/M_A$ | 2 |
| PACC2W1 | *F. pacifica* | $M_A/M_A$ | 2 |
| PACC3W1 | *F. pacifica* | $M_D/M_A$ | 1 |
| PACC4W1 | *F. pacifica* | $M_D/M_A$ | 1 |
| PACC5W1 | *F. pacifica* | $M_A/M_A$ | 2 |

| | | | |
|---|---|---|---|
| PACC6W1 | *F. pacifica* | $M_A/M_A$ | 2 |
| PACC7W1 | *F. pacifica* | $M_A/M_A$ | 2 |
| PACC8W1 | *F. pacifica* | $M_A/M_A$ | 2 |
| PACC9W1 | *F. pacifica* | $M_A/M_A$ | 2 |
| SCEC1W9 | *F. pacifica* | $M_D/M_A$ | 1 |
| SQPC17W1 | *F. pacifica* | $M_A/M_A$ | 2 |
| SQPP1W1 | *F. pacifica* | $M_A/M_A$ | 2 |
| SQPP2W1 | *F. pacifica* | $M_A/M_A$ | 2 |
| H221C4W3 | *F. transmontanis* | $M_A/M_A$ | 2 |
| H221C5W5 | *F. transmontanis* | $M_A/M_A$ | 2 |
| H221C6AW2 | *F. transmontanis* | $M_A/M_A$ | 2 |
| H2QC2W1 | *F. transmontanis* | $M_A/M_A$ | 2 |
| H2QC3W1 | *F. transmontanis* | $M_A/M_A$ | 2 |
| H2QC4W1 | *F. transmontanis* | $M_A/M_A$ | 2 |
| H2QC5W1 | *F. transmontanis* | $M_A/M_A$ | 2 |
| H2RC1W1 | *F. transmontanis* | $M_A/M_A$ | 2 |
| H2RC2W1 | *F. transmontanis* | $M_A/M_A$ | 2 |
| H2RC3W1 | *F. transmontanis* | $M_A/M_A$ | 2 |
| H2RC4W1 | *F. transmontanis* | $M_A/M_A$ | 2 |
| TELC1W9 | *F. transmontanis* | $M_A/M_A$ | 2 |
| YACC14W1 | *F. transmontanis* | $M_A/M_A$ | 2 |
| YACC4W5 | *F. transmontanis* | $M_A/M_A$ | 2 |
| YACC5W5 | *F. transmontanis* | $M_A/M_A$ | 2 |
| H221C7W9 | *F. transmontanis* | $M_A/M_A$ | 2 |
| YACC7W4 | *F. transmontanis* | $M_A/M_A$ | 2 |
| YACC8W5 | *F. transmontanis* | $M_A/M_A$ | 2 |
| BLPC2W1 | *F.* cf. *transmontanis* | $M_D/M_A$ | 1 |
| BLPP1W1 | *F.* cf. *transmontanis* | $M_D/M_A$ | 1 |
| BOBC1W1 | *F.* cf. *transmontanis* | $M_D/M_A$ | 1 |
| BOBC2W1 | *F.* cf. *transmontanis* | $M_D/M_A$ | 1 |
| BOBC3W1 | *F.* cf. *transmontanis* | $M_A/M_A$ | 2 |
| BOBC4W1 | *F.* cf. *transmontanis* | $M_D/M_A$ | 1 |
| BOBP1W1 | *F.* cf. *transmontanis* | $M_A/M_A$ | 2 |
| H1AP1W1 | *F.* cf. *transmontanis* | $M_A/M_A$ | 2 |

**Table S3.** We used RFLP genotyping to distinguish between the $M_D$ haplotype and $M_A$ haplotype in the *pactrans* complex. $M_D$ haplotype presence/absence was scored as 1/0. Colonies producing gynes (whether or not they also produced males) were assigned as 1 and colonies only producing males were assigned as 0. We collected sex ratio data at the time of collection for h221c3 and h221c7; workers from these colonies were sequenced and their supergene genotypes included in the Fisher's exact test assessing the association between the presence of an $M_D$ haplotype and gyne production.

| Colony | ID | Workers (*n*) | $M_D$ count | $M_A$ count | $M_D$ present | Gyne producing | Males collected | Gynes collected |
|---|---|---|---|---|---|---|---|---|
| 1 | ESC0028 | 5 | 0 | 5 | 0 | 0 | 11 | 0 |
| 2 | ESC0029 | 5 | 0 | 5 | 0 | 0 | 10 | 0 |
| 3 | ESC0038 | 5 | 0 | 5 | 0 | 0 | 11 | 0 |
| 4 | ESC0053 | 5 | 0 | 5 | 0 | 0 | 7 | 0 |
| 5 | ESC0054 | 5 | 0 | 5 | 0 | 0 | 13 | 0 |
| 6 | ESC0055 | 5 | 2 | 3 | 1 | 1 | 0 | 7 |
| 7 | ESC0056 | 5 | 0 | 5 | 0 | 0 | 21 | 0 |
| 8 | ESC0057 | 5 | 2 | 3 | 1 | 1 | 15 | 4 |
| 9 | ESC0058 | 5 | 0 | 5 | 0 | 0 | 10 | 0 |
| 10 | ESC0059 | 5 | 0 | 5 | 0 | 0 | 12 | 0 |
| 11 | ESC0060 | 5 | 4 | 1 | 1 | 1 | 0 | 5 |
| 12 | ESC0061 | 5 | 0 | 5 | 0 | 0 | 8 | 1 |
| 13 | ESC0063 | 5 | 0 | 5 | 0 | 0 | 13 | 0 |
| 14 | ESC0068 | 5 | 2 | 3 | 1 | 0 | 7 | 0 |
| 15 | ESC0074 | 5 | 2 | 3 | 1 | 1 | 6 | 4 |
| 16 | ESC0077 | 5 | 3 | 2 | 1 | 1 | 0 | 10 |
| 17 | ESC0092 | 5 | 2 | 3 | 1 | 1 | 0 | 17 |
| 18 | ESC0095 | 5 | 0 | 5 | 0 | 0 | 12 | 0 |
| 19 | ESC0106 | 5 | 0 | 5 | 0 | 0 | 9 | 0 |
| 20 | ESC0119 | 5 | 3 | 2 | 1 | 1 | 0 | 13 |
| 21 | ESC0134 | 5 | 0 | 5 | 0 | 0 | 15 | 0 |
| 22 | ESC0136 | 5 | 2 | 3 | 1 | 1 | 0 | 12 |
| 23 | ESC0137 | 5 | 1 | 4 | 1 | 0 | 16 | 0 |
| 24 | ESC0141 | 5 | 1 | 4 | 1 | 1 | 10 | 7 |
| 25 | ESC0143 | 5 | 2 | 3 | 1 | 1 | 2 | 3 |
| 26 | ESC0144 | 5 | 0 | 5 | 0 | 0 | 14 | 1 |
| 27 | h221c3-sequenced | 5 | 2 | 3 | 1 | 0 | male-producing | - |
| 28 | h221c7-sequenced | 5 | 2 | 3 | 1 | 1 | - | gyne-producing |
| | **Totals** | 145 | 32 | 113 | 14 | 11 | | |
